# Developing a nuclear transplantation model for Australian amphibian conservation using a common species of Australian Frog (*Limnodynastes peronii*)

**DOI:** 10.1101/2025.11.25.690599

**Authors:** Roshan Patel, Rose Upton, Simon Clulow, Brett Nixon, Bernd Gruber, Michael Archer, Michael Mahony, John Clulow

## Abstract

As amphibian populations continue to decline, there is a need to establish procedures to store and recover genetic diversity. Nuclear transplantation (NT) is a potentially useful reproductive technology with applications to conservation in specific circumstances. Although sperm have been successfully cryopreserved and used in in vitro fertilization to produce sexually mature offspring, there are many endangered and even extinct amphibian species that only exist as stored frozen adult or larval tissues. The only way to recover that genetic diversity is through NT. Nuclear transplantation can introduce new genes into captive breeding programs from individuals where no viable gametes from that individual have been cryopreserved. Amphibian NT research was common during the middle to late 20^th^ century; however, it was never conducted for amphibian conservation. Early amphibian NT experiments were conducted for the purpose of determining whether differentiation was a terminal process that involved the loss of reprogramming genes. However, following the success of mammalian NT, amphibian NT research declined, and the field has been at a standstill for decades. This study developed the first nuclear transplantation protocol for a native Australia frog using *Limnodynastes peronii*, as a model. Specifically, this paper aimed to test: 1) nuclear transplantation with fresh embryonic cells in *Limnodynastes peronii* and trial the implementation of different egg enucleation methods (pricked eggs only, UV only, and pricked + UV), 2) use of cryopreserved cells in nuclear transplantation and 3) use single nucleotide polymorphisms to determine if the nuclear genomes of nuclear transplant embryos were derived from the injected donor cell nuclei. With both fresh and cryopreserved embryonic cells as donor cell nuclei, development of the nuclear transplant embryos was low compared to the fertilized controls. Only one nuclear transplant embryo derived from a fresh embryonic cell (injected into a UV only enucleated egg) developed into a tadpole. This attained stage 36 of the Gosner Staging System for Anurans (hind limbs) before dying. As an egg activation technique, pricking the eggs resulted in higher cleavage rates and development to blastula than UV only and pricked + UV nuclear transplants (in trials with cryopreserved cells), but none of the nuclear transplants from the pricked only group developed beyond blastula. None of the nuclear transplants derived from cryopreserved embryonic cells developed to tadpoles. The most advanced stage attained from cryopreserved embryonic cells was stage 13 (neurula) from UV only eggs. Single nucleotide polymorphism (SNP) analysis testing for genotype mismatches was used to determine if the genomes of developing nuclear transplant embryos were derived from the injected donor cell nucleus. SNP analysis confirmed that two true clones were generated in this study. This is the first confirmed nuclear transplantation in any Australian native frog.

## INTRODUCTION

Amphibians are the most threatened vertebrate group in the current mass extinction. Around 41% of amphibians are at risk of extinction with 2,873 amphibian species at risk of extinction or likely extinct (Luedtke et al., 2023). Among these, 37 species are classed as extinct, and 185 species as critically endangered/possibly extinct (Luedtke et al., 2023). In Australia, 45 out of the 244 described frog species are threatened with extinction (Geyle et al., 2021) and four Australian frog species have gone extinct (Low & Booth, 2023). Some of these extinct species only have frozen tissues available. One way to reverse extinction is to utilize nuclear transplantation to reconstruct individuals of threatened and extinct species (Shapiro, 2017).

Nuclear transplantation (NT, also referred to in the literature as nuclear transfer) is a process in which individuals are produced via injecting donor cell nuclei into eggs whose maternal DNA has been removed (enucleated eggs) (Wilmut et al., 2002). The first vertebrates to be successfully cloned were *Rana pipiens* and *Xenopus laevis*, which resulted in the production of sexually mature adults of both species (Gurdon et al., 1958; McKinnell, 1962). These two frog species were the best studied species in nuclear transplantations during the mid-late 20^th^ century. Eventually, nine other amphibian species were successfully cloned (Ellinger & Carlson, 1978; Elsdale et al., 1960; Gallien et al., 1973; Gurdon, 1961; Gurdon, 1962; McKinnell, 1962; Nishioka, 1972; Ortolani et al., 1966; Signoret et al., 1962). Despite the NT protocols that existed for many amphibian species, NT experiments were not conducted for the purpose of conservation. Instead, NT experiments were used to investigate the core question in developmental biology (i.e. whether cellular differentiation was a terminal and irreversible process), and in practice to investigate if the genome of a somatic cell functionally retained all the genes represented in an organism’s entire genome (McKinnell, 1978). Since the early amphibian NT experiments, many researchers have moved to mammalian cloning models once NT was successful in mammals (Gurdon, 2013; Gurdon & Melton, 2008; McGrath & Solter, 1983). This left the field of amphibian cloning at a standstill for decades. One of the reasons mammalian NT replaced amphibian NT is that mammalian models such as mice are considered better models for human biology (Blum & Ott, 2019). Successful mammalian NT provides a more similar model than amphibian NT for human cloning for the generation of stem cells in medicine and other applications (Gurdon, 2013; Gurdon & Melton, 2008).

Cloning (in this study, the term is only used to refer to nuclear transfer/transplantation) can help in the conservation of amphibians by recovering lost genetic diversity from frozen materials such as somatic and/or embryonic cells. In this context, it also opens the possibility of recovering individuals from extinct and/or endangered species that only have frozen tissues available and no stored gametes (Herrick, 2019; Kouba & Vance, 2009; Wisely et al., 2015; Zimkus et al., 2018). Cell nuclei from those frozen tissues may be transplanted into the enucleated eggs of a common relative through interspecies nuclear transfer (iNT) (Clulow et al., 1999; Mrowiec et al., 2021). The reconstructed individuals might then be incorporated into captive breeding program (e.g. the non-amphibian examples, black-footed ferrets and Przewalski’s horse that were produced via interspecies NT were derived from cryopreserved cells of individuals that have no living descendants or were underutilized in captive breeding) (Novak et al., 2024; Novak, Ryder, et al., 2025). Nuclear transplantation brought back lost genetic diversity that can be used to counter inbreeding depression in captive breeding programs for black-footed ferrets and the Przewalski’s horse (Novak et al., 2024; Novak, Ryder, et al., 2025). This shows that and iNT can be used to introduce new founders to captive breeding programs, and recover otherwise lost genetic diversity (Wisely et al., 2015). While iNT has been successful in endangered mammal conservation, there are no cases of NT or iNT being applied successfully for amphibian conservation, nor any studies that reported the production of nuclear transplant amphibians from cryopreserved cells. Having NT and iNT as a strategy for amphibian conservation will allow in the future for frozen tissues and cells to be used to reconstruct individuals which can allow the genetic diversity from those frozen samples to be restored (Clulow et al., 1999; Novak, Brand, et al., 2025), including potentially in cases such as the gastric brooding frog (*Rheobatrachus silus*) where there are no living animals.

In this study, we report attempts to create a viable nuclear transplantation model for amphibian conservation using *Limnodynastes peronii*, a common Australian ground frog. The paper aims to: 1. Attempt NT using fresh embryonic cells and determine whether egg various enucleation procedures prior to injection affect developmental outcome, 2. Use cryopreserved embryonic cells in NT, and 3. Use genotyping to demonstrate that the nuclear transplants are true clones and only have the DNA of the donor cell nuclear genome through single nucleotide polymorphism (SNP) analysis.

## MATERIALS AND METHODS

### Animal Husbandry

Male and female *L. peronii* were collected opportunistically from Kooragang Island, Watagan National Park, and Seal Rocks, NSW opportunistically and housed at the University of Newcastle (see supplementary table 1). Individuals were housed in plastic terraria (30 ×18 x 20 cm), with 75% terrestrial environment (autoclaved gravel, leaf litter, and fake plants) and 25% aquatic environment (aged tap water). Each terrarium had one male and two females. Frogs were fed live crickets coated with calcium and vitamin powder (Multical Dust; Vetafarm, Wagga Wagga, NSW, Australia) and water was changed twice a week. All individuals underwent a prophylactic heat treatment protocol to eliminate chytridiomycosis. The frogs were placed in UV- and temperature-controlled cabinets (TRISL – 1175, Thermoline Scientific Equipment, Wetherill Park, NSW) on a 14:10 UV light cycle starting at 24°C. Temperature was increased at 2°C per day until 36°C was reached, when the temperature was changed to 37°C for six hours. Thirty-seven degrees Celsius has been shown to kill *Batrachochytrium dendrobatidis* within four hours (Johnson et al., 2003). Subsequently, temperature was reduced at 2°C per day until the original 24°C was reached. Six male *L. peronii* and 11 female *L. peronii* were used in this study.

All experiments were carried out in accordance with the University of Newcastle’s ethics approval, A-2022-207. Animals were collected and held under NSW Scientific Licence SL101269.

### Micropipette forging

Micropipettes for nuclear transplantation were constructed using a needle puller (Stutter Instruments Model P-1000) (program: heat: 499, pull: 120, Time: 100, Pressure: 200), microforge (Narishige MF-900), and a bevelling wheel (Kyowa Tokyo). Glass capillaries were manufactured by Clark Electromedical Instruments, UK. All micropipettes that were used for *L. peronii* nuclear transplantations had an inner diameter between 20-25 µm for injections and less than 10 µm diameter for initial pricking (see below). Micropipettes were sterilized in a Class II biological safety cabinet (Gelaire) with 45 minutes of UV light prior to use in NT experiments.

### Controls for nuclear transplantation

Before separating amplecting pairs to obtain unfertilised eggs for nuclear transplantation, pairs of *L. peronii* adults were allowed to produce fertilized spawn to serve as controls for scoring egg quality. Development of the natural spawn was recorded to confirm egg quality. Negative controls for NT experiments were generated by of pricking eggs at the animal pole.

### Embryo Collection

*L. peronii* spawn (n=6) were collected from spawning pairs to provide donor cells (fresh and/or cryopreserved) for nuclear transplantations. Spawns were held in 150 x15 mm petri dishes containing 10% Niu-Twitty solution (McKinnell, 1978). Spawns were kept at 13°C (Herbert, 2004) overnight in a temperature cabinet (Thermoline Scientific model: TRH-300-GD) to slow down development to synchronise Gosner stage development of donor embryos with the availability of freshly ovulated eggs. Embryos from the following embryonic Gosner stages were used to obtain donor cells: blastula (stage 9) and gastrula (stage 10-12) (Gosner, 1960).

### Cell preparation

#### Preparation of Fresh Donor Cells

Stage 9-10 *L. peronii* embryos (Gosner, 1960) were dejellied using 1% cysteine for 2 minutes (Sigma-Aldrich cat 1276) (prepared in calcium and magnesium free Niu Twitty (also known as modified Niu Twitty) ((50.36 mM NaCl, 0.67 mM KCl, 3.65 mM Na_2_HPO_4_, 0.85 mM KH_2_PO_4_, 2.38 mM NaHCO_3_; pH: 7.8)) (McKinnell, 1978) and pH adjusted to 8.0 using NaOH and 0.22 µm filtered)). Dejellied embryos were washed in 70 µg/mL gentamicin sulphate (Indofine 1405-41-0) made up in calcium and magnesium free Niu Twitty (0.22 µm filtered) for ten minutes. Donor cells were dissociated via dissection of single (n=2) or a mixture of 15 embryos (n=1) with forceps and/or glass needles. Dissection was completed in 2.5% w/v Bovine serum albumin (BSA) (Sigma-Aldrich A7906) + 0.5 mM ethylenediaminetetraacetic acid (EDTA) (Sigma: E-5134) made up in calcium and magnesium free Niu Twitty (and 0.2 micron filtered). The dissociated cells were washed with 70 µg/mL gentamicin for 10 minutes and the gentamicin was washed out via dilution with 5% w/v BSA (made in calcium and magnesium free Niu-Twitty and 0.22 micron filtered).

#### Preparation of Frozen Donor Cells

*L. peronii* gastrula cells (Gosner stages 10-12) from four spawns were cryopreserved with 10% v/v dimethyl sulfoxide (DMSO) (Ajax Finchem: AJA2225) + 10 % w/v sucrose (Ajax Finechem: AJA530) (10D10S). Gastrula cells were cryopreserved and used in NT instead of blastula cells due to gastrula cells having better post-thaw recovery than blastula cells. Gastrula cells from four different spawning pairs were cryopreserved. Either one embryo (dissociated in 300 µl of 2.5% BSA; three embryos dissociated separately) (n=1) or a pool of 30 embryos (n=3) were dissociated (in 2 mL of 2.5% BSA) and cryopreserved. From one spawning pair, three gastrula embryos were dissociated individually while 30 gastrulas from each of the other three spawn were dissociated to provide cells for cryopreservation. Prior to cryopreservation, 250 µl aliquots of cryoprotectant were added to 50 µl aliquots of embryonic cell preparations and placed in Eppendorf tubes on ice. 300 µl of the combined cryoprotectant and cell mixtures were loaded into 500 µl straws (Minitube EcoStraw, cat 13408/3010) and sealed with polyvinyl alcohol powder. At least three straws were prepared from each spawn. Straws were loaded into a programmable freezer (Cryologic, Australia, Model: CL-3300) and the freezing protocol used was as per (Lawson et al., 2013). Briefly, straws were held at 10°C for ten minutes, cooled to −7°C at a rate of −1°C per minute, held at −7°C for 10 minutes, cooled to −30°C at −1°C per minute, held at −30 °C for 10 minutes, and then the temperature was allowed to free fall from −30°C until −80°C was reached. Straws were then plunged in liquid nitrogen and transferred to a dewar to be stored.

Four nuclear transplantation trials (with cryopreserved cells) were completed with single dissociated embryos, and four NT trials were undertaken with multiple dissociated embryos. Straws were stored in liquid nitrogen and were thawed whenever eggs became available for NT experiments. Straws were thawed at room temperature into a 5 cm petri dish and 50 µl of the 300 µl cell + cryoprotectant solution was taken for post-thaw counts (membrane integrity and cell concentration, see supplementary Figure 1). Cryoprotectant was removed via dilution with calcium and magnesium free Niu Twitty. This was achieved by pipetting 9 ml of calcium and magnesium free Niu Twitty slowly pipetted into the dish containing the thawed cells. Then, 6 mL of the cell/cryoprotectant solution was pipetted out and 1 mL of 5% BSA was added to the remaining 3 mL of the cell/cryoprotectant mixture. The estimated concentration of residual cryoprotectant after the dilution was 0.207%. Cryoprotectant was diluted out instead of spinning the cells down (washing the cells) to limit physical damage. Cells were washed in 70 µg/mL gentamicin prepared in calcium and magnesium free Niu-Twitty for 10 min. The gentamicin was diluted out with calcium and magnesium free Niu-Twitty and 5% BSA (made in calcium and magnesium free Niu-Twitty). Cells were kept in 5% BSA during the transplantation.

#### Cell counts and scores (membrane integrity and concentration) for fresh and thawed cell preparations

Following the above cell preparation methods, embryonic cells were examined in a haemocytometer under a Nikon eclipse e200 microscope at 400x magnification for cell membrane integrity staining and concentration (cells/mL) counts. Two replicate counts were completed per sample. Ten microlitres of undiluted cell suspension was pipetted into each chamber and cells were counted in 4-9 primary squares to determine concentration. A trypan blue stain exclusion assay was used to assess cell membrane integrity. Cell suspensions were stained with 0.5% trypan blue at a 1:1 ratio, with a volume of 10 µl viewed at 400x magnification in a haemocytometer. Unstained cells were counted as plasma membrane intact (PMI) and cells that stained blue were score as not intact. A target of 100 cells per count was made; however, observing a minimum of 100 cells per count was not always possible in cryopreserved samples due to the low number of cells observed post-thaw. Pre-freeze assessment was completed for the cryopreserved cells using the same methods.

### Egg collection and preparation

Eggs from *L. peronii* were collected by separating amplecting pairs that were in the process of spawning. Where ovulating females were observed at night, females were placed at 10°C overnight to delay egg release until cells for microinjection could be prepared the following day. Eggs were stripped from an ovulating female *L. peronii* via gentle massaging of the abdominal area into a petri dish fill with 100% Niu Twitty solution (58 mM NaCl, 0.67 mM KCl, 0.3 mM Ca (NO_3_)_2_, 0.41 mM MgSO_4_*7H_2_O, 0.31 mM Na_2_HPO_4_, 0.15 mM KH_2_PO_4_, 2.38 mM NaHCO_3_; pH: 7.8) (0.2 µm filtered) (McKinnell, 1978). Female *L. peronii* that were kept at 10°C overnight were brought to room temperature before egg collection. Eggs were maintained in 100% Niu Twitty for at least 15 minutes before enucleation and/or activation. In *X. laevis*, eggs undergo contraction at the animal pole 10-20 minutes after laying and this offers increased resistance to physical stress such as penetration of the micropipette compared to eggs that have not undergone contraction (Gurdon, 1960).

For NT trials using fresh embryonic cells, eggs from two different female *L. peronii* and donor cells from two different spawning pairs were used in three different trials. One female and one spawning pair provided eggs and donor cells on two different occasions. For NT using cryopreserved embryonic cells, eggs from five different female *L. peronii* and cryopreserved embryonic cells from four different spawning pairs were used for eight trials. One spawning pair provided cells four times, and another provided donor cells twice (see Table 1).

**Table 1:** The experimental configuration for nuclear transplantation trials (egg and cell donor IDs, enucleation treatments and cryopreservation).

| trial | Aim | egg donor ID | donor cell ID | stage donor cell | donor cell cryopreserved | enucleation treatment<sup>1</sup> | eggs injected |
| --- | --- | --- | --- | --- | --- | --- | --- |
| 1 | 1 | female 1 | spawning pair 1 | stage 9/10 | no | UV only | 39 |
| 2 | 2 | female 2 | spawning pair 2 | stage 10/11 | yes | UV only | 63 |
| 3 | 2 | female 3 | spawning pair 3 | stage 10 | yes | UV only | 56 |
| 4 | 1 | female 1 | spawning pair 4 | stage 10 | no | UV only | 30 |
| 5 | 2 | female 2 | spawning pair 5 | stage 10 | yes | UV only | 57 |
| 6 | 2 | female 4 | spawning pair 5 | stage 10 | yes | UV only | 72 |
| 7 | 2 | female 4 | spawning pair 2 | stage 10/11 | yes | UV only | 12 |
|  |  |  |  |  |  | Pricked + UV | 22 |
|  |  |  |  |  |  | Pricked only | 12 |
| 8 | 2 | female 3 | spawning pair 5 | stage 10 | yes | UV only | 17 |
|  |  |  |  |  |  | Pricked + UV | 20 |
|  |  |  |  |  |  | Pricked only | 20 |
| 9 | 2 | female 6 | spawning pair 5 | stage 10 | yes | UV only | 29 |
|  |  |  |  |  |  | Pricked + UV | 30 |
|  |  |  |  |  |  | Pricked only | 10 |
| 10 | 2 | female 5 | spawning pair 6 | stage 12 | yes | UV only | 20 |
|  |  |  |  |  |  | Pricked + UV | 23 |
|  |  |  |  |  |  | Pricked only | 25 |
| 11 | 1 | female 4 | spawning pair 1 | stage 9 | no | UV only | 18 |
|  |  |  |  |  |  | Pricked + UV | 28 |
|  |  |  |  |  |  | Pricked only | 30 |

<sup>1</sup> Most nuclear transplantation trials were initially performed with UV only eggs. Pricked only and pricked + UV egg treatments were introduced at trial 7.

Recipient eggs were washed in 10% v/v Niu Twitty (0.2 µm filtered) and placed on small pieces of filter paper (animal pole facing up) using forceps (this allowed eggs to be easily moved via forceps). Eggs were prepared by one of three enucleation methods prior to injection:

- eggs were placed in a Uvitech chamber (model number: CL-508) and exposed to UV for 42 seconds to enucleate them (Gurdon, 1960);
- eggs were pricked with a glass micropipette in the animal pole (Briggs & King, 1952) and then enucleated with UV for 42 seconds;
- eggs were pricked and not enucleated. Enucleated eggs were injected right away or kept in 100% Niu Twitty at room temperature to be injected later (within five hours). Eggs that were only pricked, but not enucleated, were injected right away before the egg cleaved.

### Nuclear transplantation procedure

Nuclear transplantations were performed opportunistically whenever ovulated eggs became available. The method for injecting cells into eggs was adapted from the protocols for *R. pipiens* and *X. laevis* (Briggs & King, 1952; Elsdale et al., 1960). Prior to picking up cells, insides of the micropipettes were coated with 7% polyvinylpyrrolidone via aspiration (Sigma: P-5288, made up in calcium and magnesium free Niu-Twitty and 0.22 µm filter sterilized) to prevent cells from sticking to the micropipette and to allow proper fluid control within the micropipette during nuclear injection (Geber et al., 2002; Hiraoka et al., 2021; Kato & Nagao, 2012). Donor cells were maintained either briefly on ice (approximately 5-10 mins) until nuclear transplantation (one trial for fresh and four trials for cryopreserved) or at room temperature (two trials for fresh and four trials cryopreserved). Cells with larger diameters than the micropipette’s inner diameter were aspirated to rupture the plasma membrane, so when the donor cell nucleus was injected into the egg, it was in contact with the egg’s cytoplasm. Eggs (UV only, pricked + UV, or pricked only (still attached to filter paper)) were taken out of the 100% Niu-Twitty and placed in a dry petri dish. The broken cell was injected into the animal pole, and the egg was quickly placed into 100% Niu-Twitty for at least an hour to allow the injection site to heal (Elsdale et al., 1960). Then, it was replaced with 10% Niu-Twitty. The nuclear transplantation system consisted of the following: Olympus SZH-ILLB dissecting microscope, Olympus LG-PS2 external light source, Leitz micromanipulator, Eppendorf cell tram oil 5176, and mineral oil (Sigma:400-5).

It was more challenging to conduct NT with fresh embryonic cells than cryopreserved cells as there were few occasions where both donor eggs and freshly prepared embryonic cells from fertilised embryos were available at the same time.

#### Assessment of nuclear transplant development

An hour after the last nuclear transplantation in each run, injected eggs were examined underneath an Olympus szx16 to look for any signs of development (cleavage, blastula, gastrula, neurula, and tadpole). Development was monitored daily. Nuclear transplants were kept at room temperature to develop. Photo images were taken with a Tucsen ISH500 camera under the Olympus szx16 microscope. Any tadpoles that reached feeding stage were placed in small plastic containers (14.5 cm X 9.5 cm X 7 cm) with 10% Niu-Twitty (around one litre) for rearing (eventually replaced with 10% Holtfreter’s solution). Tadpoles were fed spirulina (Nature’s Way) and trout pellet (Ridley’s Agricultural Products, NSW). Spirulina and trout pellet was given ad libitum three times a week.

### Single Nucleotide Polymorphism (SNP) preparation and analysis

Nuclear transplant embryos that displayed no abnormalities were allowed to continue developing, while abnormal/dying nuclear transplants were fixed in 80% ethanol. Dissociated donor cells and foot webbing from the adult *L. peronii* (egg and spawn donors) were fixed in 80% v/v ethanol. Samples were sent to Diverse Array Technologies, Canberra for single nucleotide polymorphism (SNP) sequencing (see below) to determine whether nuclear transplantation embryo nuclear genomes were derived from the injected donor cell or if the maternal pronucleus contributed to the nuclear genome of the developing embryo (or potentially both).

#### Statistical analysis of cell viability during development of the NT embryos

For cell membrane integrity data pre-freeze and post-thaw (for donor cells used in NT), general linear mixed modelling (GLMM) was used to fit binary logistic regressions (proportions of intact embryonic cells) with total number of cells equalling the weights for the model. Cryoprotectants were used as the main effects in the models and overdispersion was addressed using an observation level random effect. For cell concentration (cells/mL), GLMM were used to fit a zero inflated negative binomial distribution. The raw counts were used with an offset of the log reciprocal of an adjustment factor (based on dilution factor, volume and the number of squares counted in the haemocytometers) to convert the raw counts to cells/mL for data processing in the models. Cryoprotectant was used as the main effects, dissociation was the random effect, and the negative binomial distribution was used to account for overdispersion.

For NT completed with fresh embryonic cells, Generalised Linear Mixed Models (GLMM) were used to fit binary logistic regressions (interpreted as the proportion surviving embryos at cleavage, blastula, gastrula, neurula, and tadpole) with total number eggs equalling the weights for the model. Interaction of treatment (NT_UV vs natural spawn) and embryonic stage were used as the main effects and overdispersion was addressed using an observation level random effect. Methods of cell dissociation and cells on ice were tested as as random effects, but ultimately observation level random effect was chosen as it best dealt with overdispersion in the data. There was only one replicate of Prick and Prick+UV treatments, so they were not included in the model.

For the development of NT embryos from cryopreserved cells, GLMM were used to fit binary logistic regressions (interpreted as the proportion surviving embryos at cleavage, blastula, gastrula, neurula, and tadpole) with total number eggs equalling the weights for the model. Interaction of treatment (NT_UV vs NT_Prick vs NT_Prick_UV vs natural spawn) and embryonic stage were used as the main effects and overdispersion was addressed using an observation level random effect.

All data analysis was completed in the R programming language (Version 4.1.2) in RStudio with the following packages: glmmTMB was used for proportion development of NT embryos from cryopreserved cells and cell concentration (Brooks et al., 2017); lme4 for general linear mixed modelling (GLMM) of proportion data (membrane integrity and development of NT embryos from fresh embryonic cells) (Bates, 2014); emmeans was used to calculate Estimated Marginal Means (EMM), 95% confidence intervals (CI), and inverse odds ratios (OR) for cell membrane integrity, concentration (cells/mL), and proportion development (for NT embryos from fresh and cryopreserved cells) (Lenth, 2022); DHARMa was used to assess QQplot residuals, distribution, dispersion, and uniformity (Hartig, 2018); gridExtra, ggplot2 and ggeffects were used for plotting (Auguie & Antonov, 2017; Wickham et al., 2019; Wickham et al., 2024).

#### SNP Analysis

One plate of 94 samples was sent to Diversity Arrays Technologies for SNP sequencing. From the 94 samples (including 73 nuclear transplants from all three egg treatments) that were sent for DaRtseq sequencing to Diversity Arrays Technologies, 55 individual samples had DNA of sufficient quality to complete sequencing, and sequency generated data for 85,076 SNPs. The following packages were used for SNP analysis In the R programming language (4.5.0): DartRverse (version 1.0.6) and stringr (version 1.5.1).

Filtering of SNP data was performed to select high quality SNP loci for analysis. The function gl.filter.allna was used to remove loci and individuals that had missing values. The gl.filter.locmectric function was used to filter read depth for reference and alternative alleles to greater than seven and less than 100. Following read depth filtering, gl.filter.reproducibility was used to retain SNPs with reproducibility of greater than 99%. Next, the gl.filter.monomorphs function was used to detect and remove monomorphic loci, however, none were detected. Individual and loci call rate were removed using the gl.filter.callrate function with a threshold of greater than or equal to 85. Secondary SNPs were removed with the gl.filter.secondaries function and monomorphic loci were removed again with gl.filter.monomorphs. Finally, minor allele frequency (alleles with low frequency) was removed from the data, with the gl.report.maf function with a threshold of 0.0132. After filtering, 38 individual samples (17 nuclear transplants and the rest samples from the donor cell DNA or egg donor) remained with 1108 SNPs remaining after the SNP data filtering regime had been applied.

Once filtering of SNP data was completed, genotype mismatches and exclusions were calculated between nuclear transplant genotypes and their parent (cell nuclei donors) genotypes. Mismatches are identified, for example, if both parents (donor cell pair) have an AA genotype for a locus, and a tested nuclear transplant genotype is B, then the nuclear transplant genotype is a mismatch and cannot be derived from the parents (See Table 2 for example). Mismatch checks were based on allele combinations of the nuclear transplant and the two donor parents that were impossible through mendelian inheritance. Once mismatches were counted across all the SNPs analysed for each individual NT cell donor, the egg donor genotypes were assessed to determine if they could explain the number of mismatches in the SNP datasets. For each mismatch that was detected between the nuclear transplant and the two donor parents at a specific locus, the egg donor’s genotype was compared to those mismatches. If there was a mismatch between the NT genotype and the egg donor’s genotype, it was counted as an exclusion (i.e. for that specific SNP locus).

**Table 2:** Rationale for interpretation of mismatch data and exclusion for donor parents, nuclear transplant, and egg donor.

| Parent 1 Genotype | Parent 2 Genotype | Nuclear Transplant Genotype | Mismatch | Egg Donor Genotype | Exclusion From analysis | True nuclear clone |
| --- | --- | --- | --- | --- | --- | --- |
| AA | AA | AA | no | BB |  | Possible |
| AA | AA | AB | yes | BB | no | No |
| BB | BB | AB | no | AA | yes | Uncertain |

The mismatch and exclusion values were summed up for each nuclear transplant and its corresponding parents across all 1108 SNPs (it is necessary to do this across a large number of SNP as genotyping errors can occur in the sequencing data, and must be excluded statistically). In addition, the number of genotype mismatches relative to all possible donor parent pairs were calculated, which provided a distribution of mismatch counts for each nuclear transplant against the background of all the donor parent pairs (the quantile). Mismatches and exclusions were calculated for nuclear transplants that had both donor parents (donor cells were excluded). The underlying rationale for interpreting mismatch occurrences is the following: mismatches should not happen (if NT nuclear genome is the same as the donor cell nuclear genome) but may occur if egg donor DNA has been incorporated into the developing embryonic NT genome.

After mismatches and exclusions were counted, error rates in the genotyping were estimated from pairs of replicated genotype likelihoods by establishing a confusion matrix to record how often a pair of genotypes occurred. Then, the observed genotypes were compared to the expected genotypes and a log-likelihood function was used to calculate the error rate.

Error rates were calculated for nuclear transplant pairs from each trial using the data for all trials by summing data across all genotypes. This allowed the average genotyping error rate to be calculated across the entire SNP data set. Trials with only one nuclear transplant were left out of the error rate calculation.

The average genotyping error rate was used to check how many mismatches would occur if applied to the parents and the offsprings by running a genotyping error (mismatch) simulation. This was accomplished by running 500 simulations, in which the estimated error rate was introduced to the parents and nuclear transplant data. The expected number of mismatches generated for the estimated error rate were calculated based on the average error rate and average genotype frequencies (calculated for all the nuclear transplants from the filtered dataset). The quantile function was used to examine the distributions of the mismatches for the 2.5%, 50%, and 97.5% quantiles. Where the observed mismatches were less than would have been expected with the estimated error rate, it was accepted as strong evidence that the nuclear transplant is the result of a mendelian pairing of the parents (i.e. a true clone).

## RESULTS

### Membrane integrity and concentration of the donor cells (cryopreserved and non-cryopreserved)

As an indicator of donor cell quality, the plasma membrane integrity of donor was assessed (Supplementary Figure 1).

There was a significant effect of cryopreservation on membrane integrity (likelihood ratio test (LRT) χ^2^ (1) = 51.793, P= 6.168X10^-13^) and concentration (likelihood ratio test (LRT) χ^2^ (1) = 11.347, P= 0.00076) of gastrula cells pre-freeze and post-thaw (cryopreserved with 10D10S) (See supplementary Figure 1). Odds ratios and 95% confidence intervals (CI) were generated to compare the effect of cryopreservation on membrane integrity and concentration. Pre-freeze membrane integrity (EMM: 98.9%) and concentration (EMM: 2.9 x 10^5^ cell/mL) were significantly higher compared to the post-thaw membrane integrity (EMM: 69%, OR: 40.3, 95% CI: 20.116-80.728) and concentration (EMM: 112029 cells/mL, OR: 2.568, 95% CI: 1.692-3.897). For fresh embryonic cells used in NT, average percentage of membrane intact cells and concentration (cells/mL) were 97.21% and 130000 cells/mL, respectively.

### Negative controls

Around 45.5% of the pricked eggs (15/33 eggs across all trials) cleaved (first cleavage only); however, the pricked eggs arrested overnight and never reached beyond Gosner Stage 2 (2 cell stage).

### The development of nuclear transplant embryos derived from fresh embryonic cells compared to natural spawn

The percentage of nuclear transplant embryos developing from first cleavage to tadpole progressively declined in comparison to natural spawned controls (Figure 1). Nevertheless, a small number of embryos were confirmed to be true clones (see SNP analysis), and the most advanced true clone reached a tadpole forming hindlimbs.

**Figure 1:**
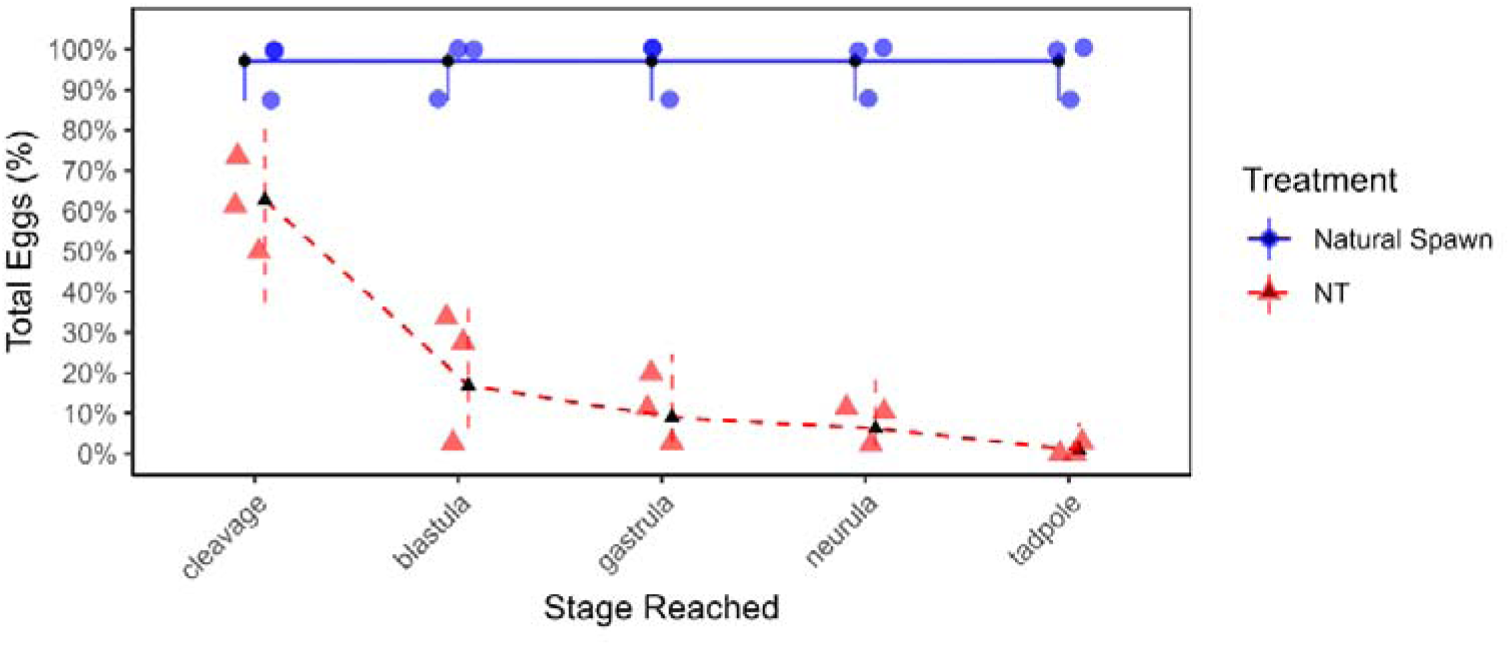
Development of nuclear transplants ((UV eggs only and natural spawn (n=3)) from fresh embryonic cells from first cleavage to tadpole stage. Estimates generated from GLMM binomial; black dots are the EMMs, error bars are 95% CI, and the raw data are the red triangles (NT) and blue dots (natural spawn).

There was a significant effect of treatment (NT vs. natural spawn) and stage interaction on the proportion of embryos surviving (likelihood ratio test (LRT) χ^2^(4) = 9.7331, P= 0.04517) with NT embryos showing decreased survival compared to embryos from natural spawn (see Figure 1).

Odds ratios and 95% CI were calculated to compare development between treatments (natural spawn versus NT with fresh (non-cryopreserved) cells. Cleavage was significantly higher in embryos from natural spawns (EMM: 97.13%) compared to NT (EMM: 62.73%), with an odds ratio of 20.14 (95% CI: 2.991-135.651). The percentage of embryos developing to blastula was significantly higher from natural spawns (EMM: 97.14%) compared to NT (EMM: 16.9%), with an odds ratio of 167.2 (95% CI: 23.168-1206.642). The percentage of embryos reaching gastrula was significantly higher in natural spawn (EMM: 97.13%) compared to NT (EMM: 9.03%), with an odds ratio of 341.1 (95% CI: 44.843-2594.93).The percentage of embryos reaching neurula was significantly higher in natural spawn (EMM: 97.13%) compared to NT (EMM: 6.3%), with an odds ratio of 505.1 (95% CI: 64.107-3979.178). The percentage of embryos reaching tadpole was significantly higher in natural spawn (EMM: 97.14%) compared to NT (EMM: 0.84%), with an odds ratio of 4014.7 (95% CI: 237.391-67894.658).

Between consecutive stages for NT embryos, there was a significant drop in the percentage of embryos surviving between cleavage and blastula (OR: 0.12 and 95% CI: 0.026-0.551), but not between blastula and gastrula, gastrula and neurula, and neurula and tadpole (see supplementary table 1). A single tadpole (figure 2C and 2D) derived from fresh embryonic cells started feeding and developed hind limbs (stage 36 of Gosner stage) before dying (Figure 3).

**Figure 2:**
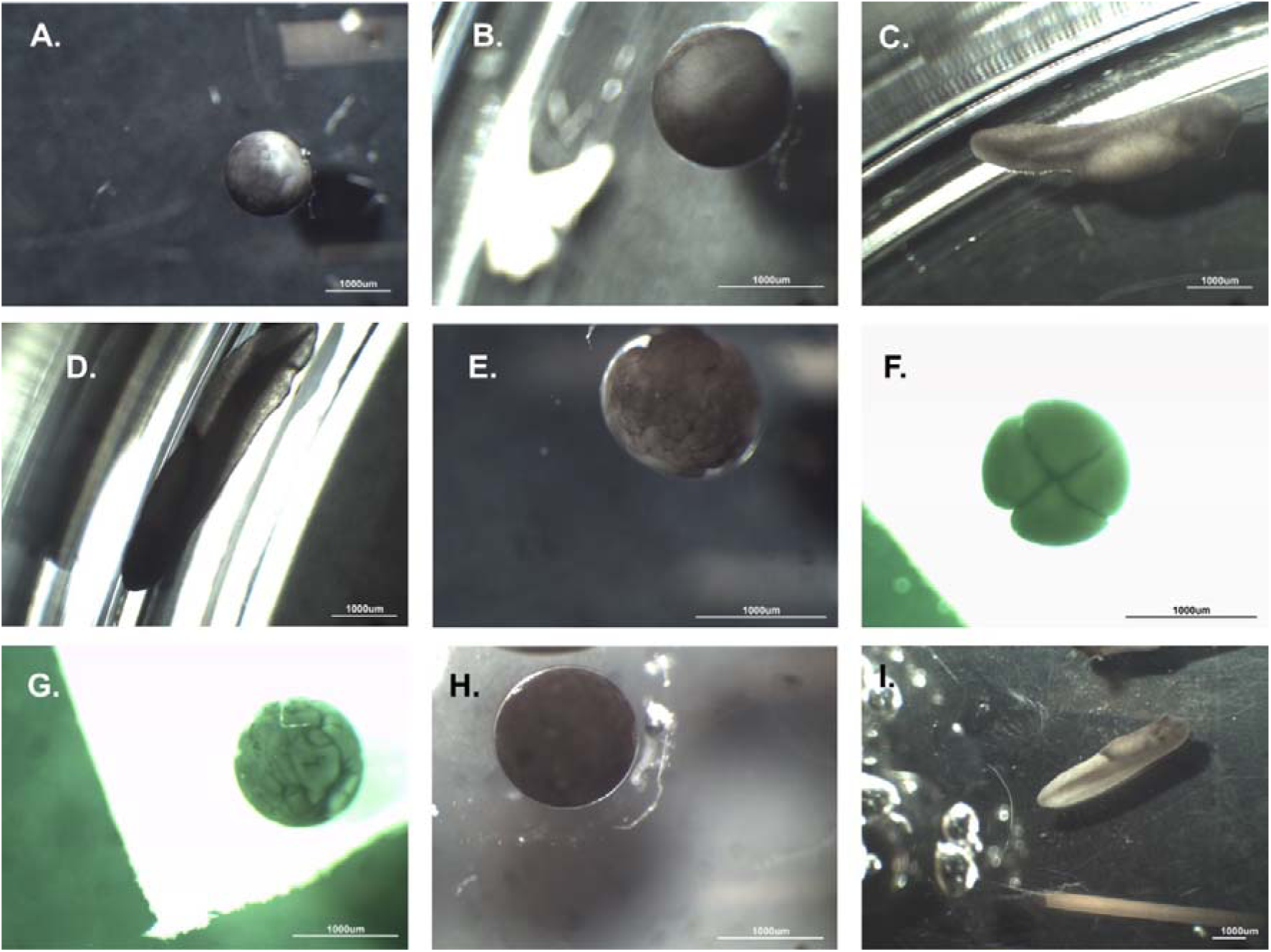
Nuclear transplants derived from fresh embryonic cells (panels 2A-2G) and natural spawn controls (panels 2H-2I). All the nuclear transplants are derived from eggs that have been exposed to UV only (no prick). 2G is a nuclear transplant that had undergone abnormal development. The nuclear transplant tadpole (2C and 2D: same individual) developed to stage 36 of the Gosner stage before dying. This tadpole was confirmed to be a true genomic clone and was the most advanced NT produced in the study.

**Figure 3:**
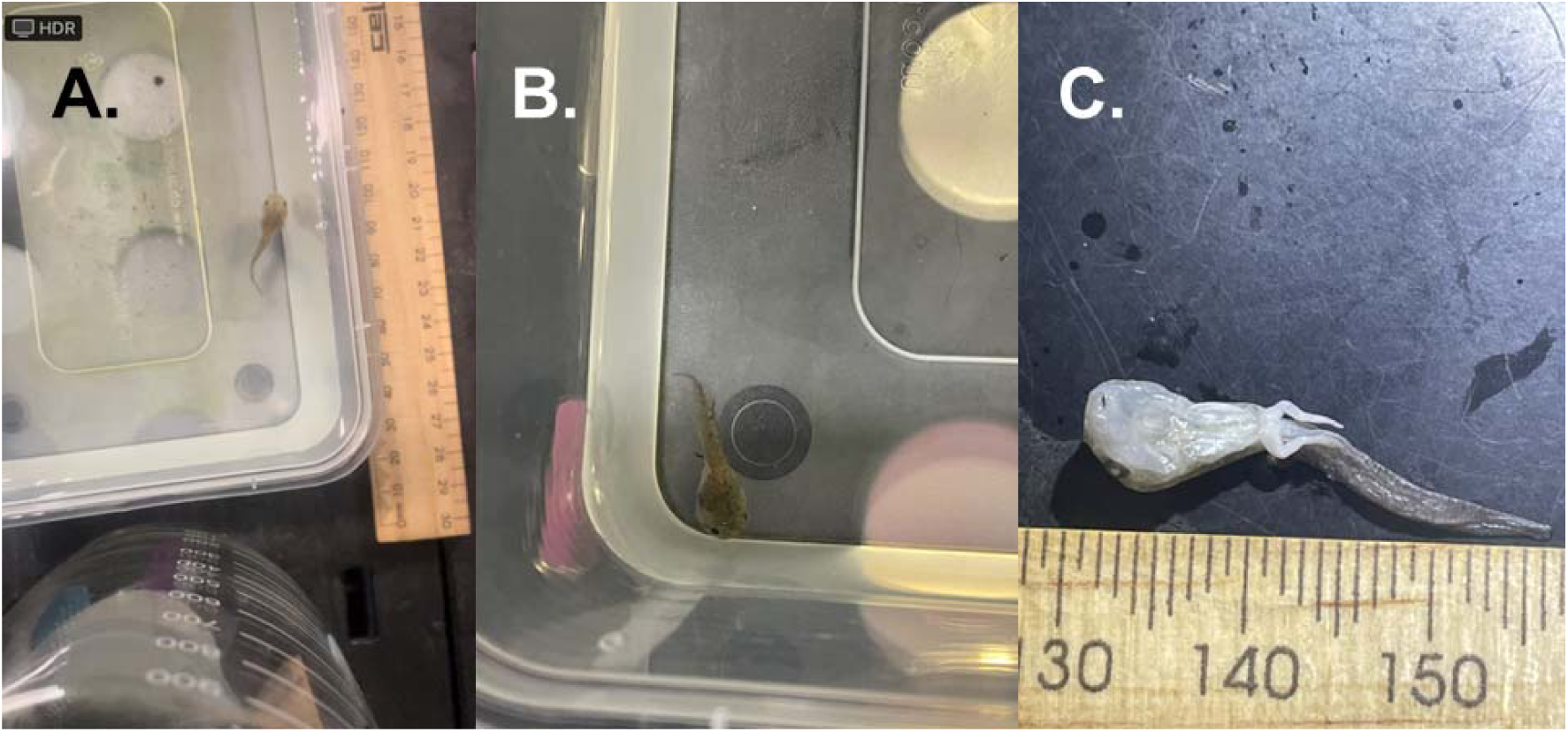
The nuclear transplant *L. peronii* tadpole derived from a fresh embryonic cell (confirmed to be true clone). 3A and 3B are the live tadpole. 3C image taken after the NT tadpole died.

There was only one trial replicate for Prick and Prick+UV treatments for NT with fresh embryonic cells (compared to three replicates for UV treatment), so could not be statistically analysed. Nevertheless, development to the neurula stage (Gosner stage 14) occurred in 2 out of 28 injected eggs in the Prick+UV eggs, and a single tadpole (stage 21 of the gosner stage) was produced from pricked only eggs (one out of 30 injected eggs). For Prick+UV treatment, the total percentage of eggs injected reaching cleavage, blastula, gastrula, neurula, and tadpole declined steadily from were 60.7%, 39.3%, 10.7%, 7.1%, and 0%, respectively. For Prick, development (percentage of total eggs injected) from cleavage, blastula, gastrula, neurula, and tadpole in the following order were 90.0%, 33.0%, 10.0%, 10.0%, and 3.3% respectively. One tadpole from the pricked group hatched and started swimming (See Figure 4C), but developed oedema and died.

**Figure 4:**
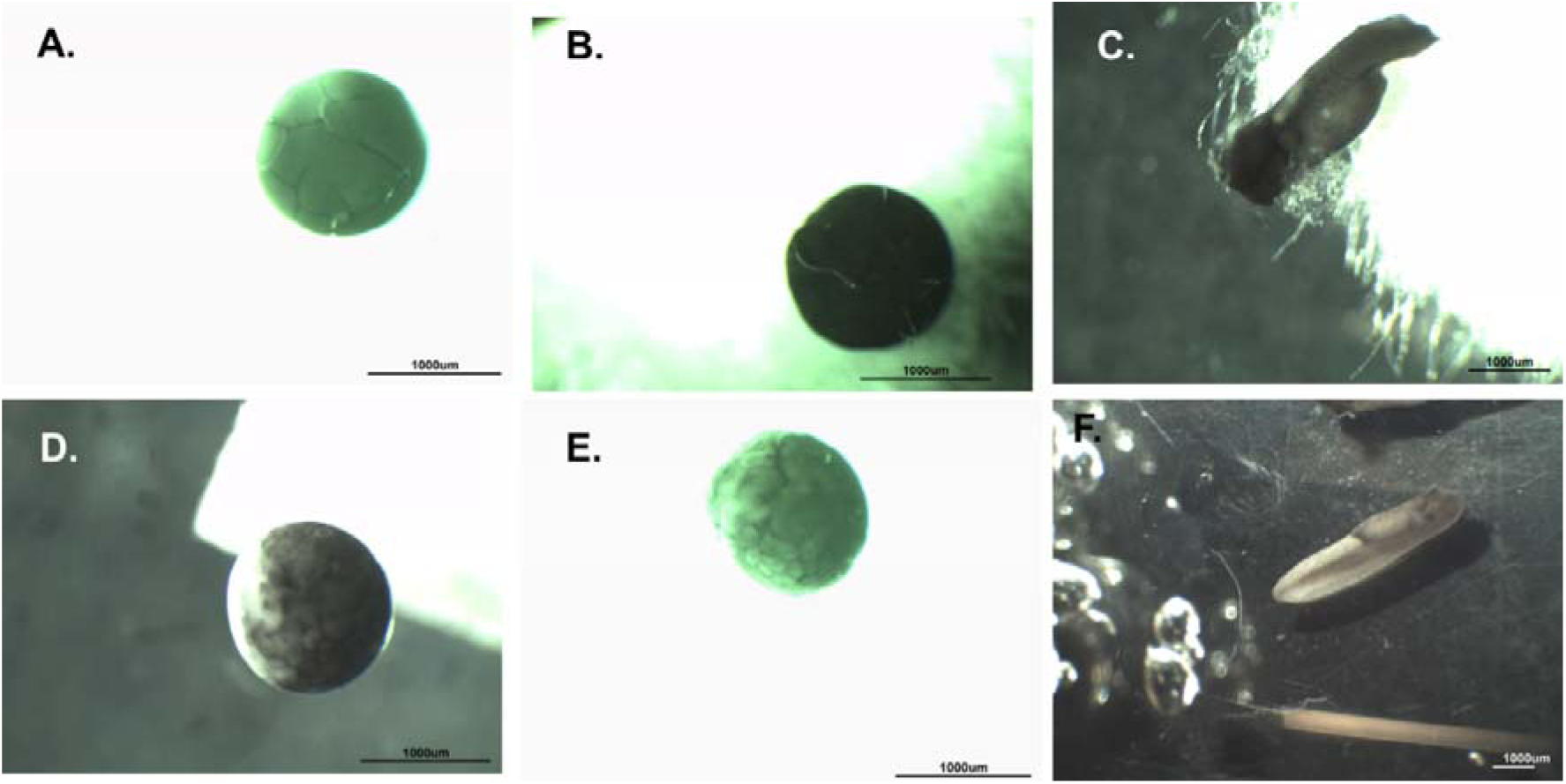
Various nuclear transplants derived from fresh embryonic cells injected into eggs that were from treatments Pricked (4A-4C) or Pricked+UV (4D and 4E). Tadpole in 4C hatched but succumbed to oedema. 4F is a tadpole from natural fertilization (spawned).

**Figure 5:**
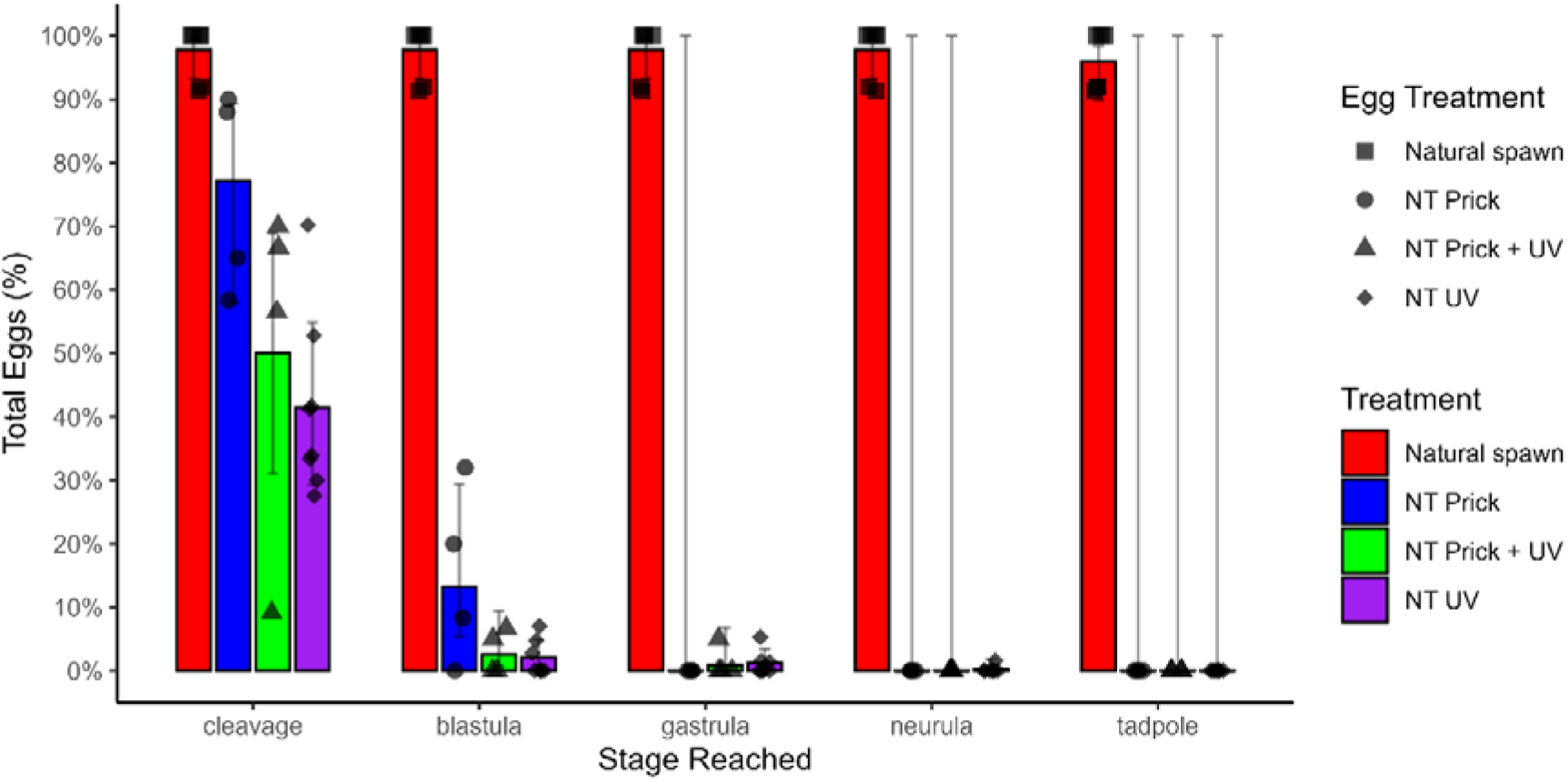
Development of nuclear transplants from cryopreserved embryonic cells (for Prick, Prick + UV, and UV treatments) and natural spawn. GLMM binomial; black dots represent estimated marginal means (EMMs), error bars represent 95% CI, and the raw data are the black shapes. Large 95% CIs development at gastrula, neurula, and tadpole stages are statistical idiosyncrasies occurring due to zero occurrences (zero embryo survival) at those stages.

### The development of nuclear transplant embryos derived from frozen embryonic cells compared to natural spawn

The most advanced stage attained for a NT embryo derived from a cryopreserved cell was a single, abnormal neurula (from UV only eggs) before arresting (figure 6E). Abnormal NT embryos were observed in all egg treatments (See figures 6 and 7). Thus, the ultimate stage of development reached in NT’s with cryopreserved cells in this study was substantially less (Gosner Stage 13, neurula) than that recorded for NT’s with non-cryopreserved cells (tadpole with hindlimbs, Gosner Stage 36).

**Figure 6:**
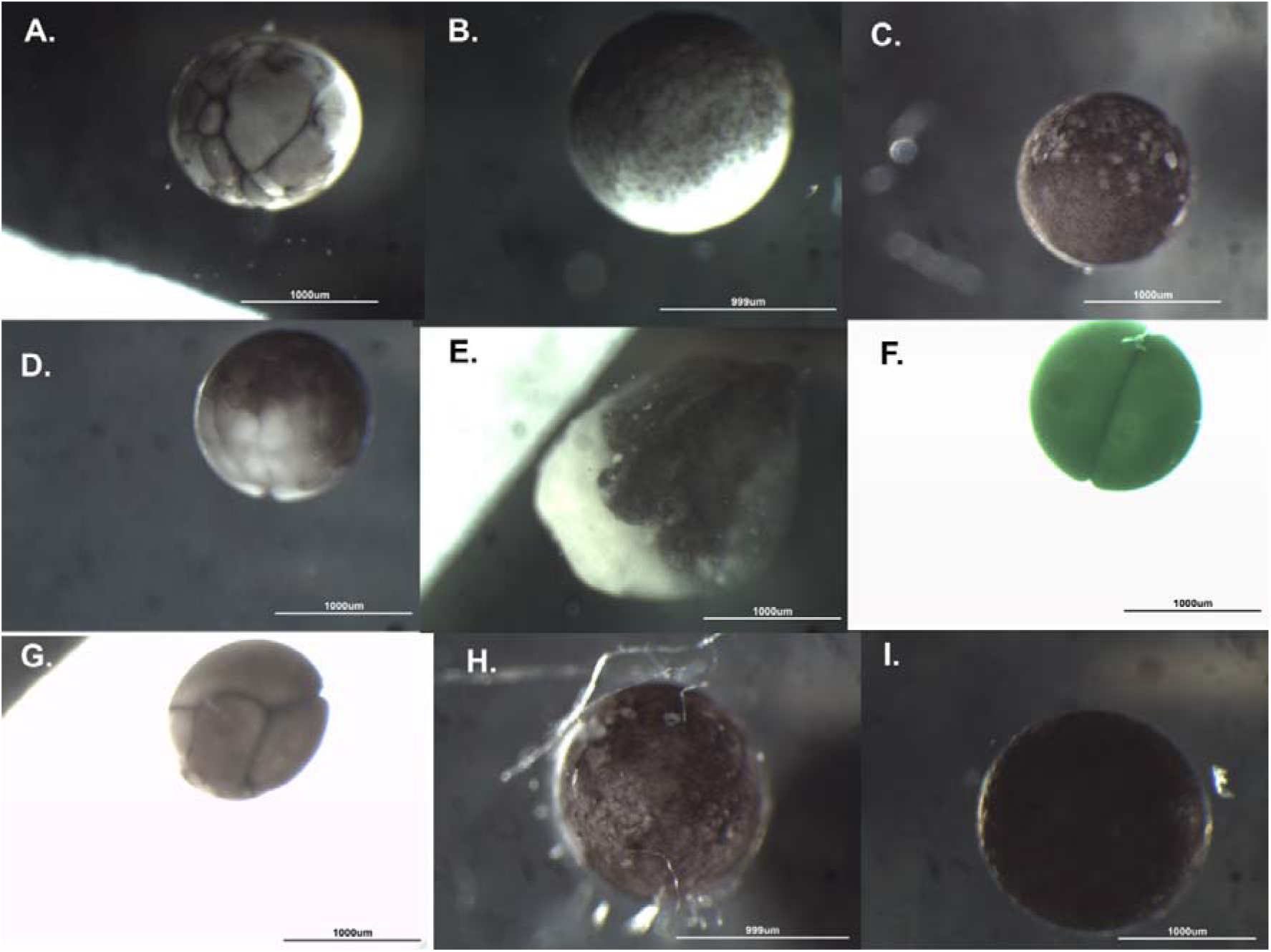
Nuclear transplants derived from cryopreserved embryonic cells (6A-6H) and compared to natural spawn embryo (6I, Gosner stage 11/12, late gastrula). 6B, 6D, and 6F are nuclear transplants that are developing normally while 6A, 6C, 6E, 6G, and 6H undergone abnormal development. The most advanced developmental stage reached with cryopreserved cells was 6E (stage 13, neurula). All the nuclear transplants are derived from eggs that have been exposed to UV.

**Figure 7:**
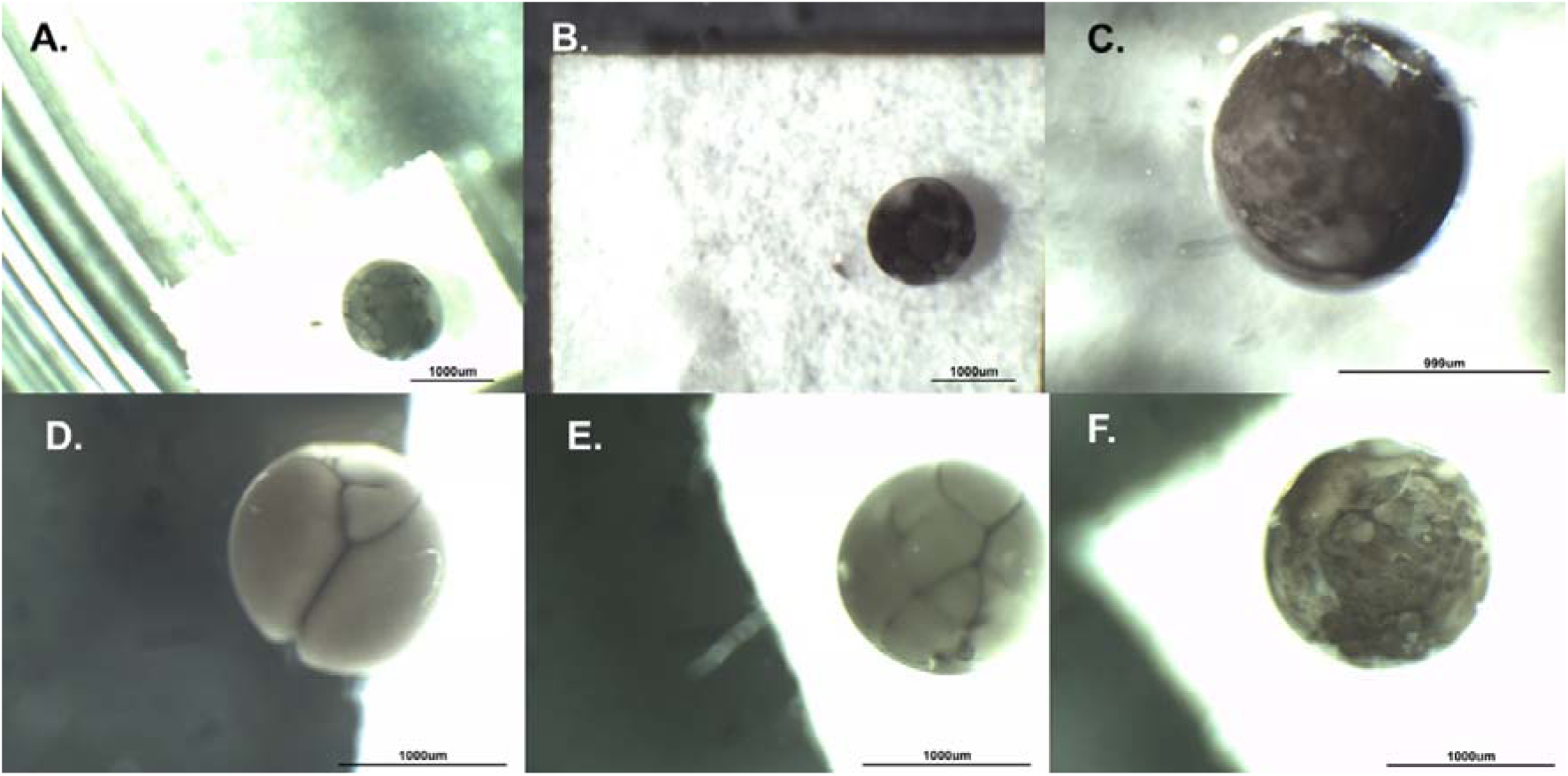
Nuclear transplants embryos derived from injecting cryopreserved cells into eggs that were Pricked+UV (7A-7C) or Pricked (7D-7F). All embryos underwent abnormal development.

There was a significant effect of treatment (UV vs. Prick vs. Prick+UV vs. natural spawn) and stage interaction on the proportion of embryos surviving (likelihood ratio test (LRT) χ^2^(12) = 43.731, P= 1.697×10^-05^) with NT embryos from all three egg enucleation treatments having decreased survival compared to embryos from natural spawn (See Figure 5).

Odds ratios and 95% CI were calculated to compare development between treatments. Natural spawns had the highest percentage of cleaving embryos (EMM: 97.798%) compared to pricked only eggs (EMM: 77.197%;13.12-fold; 95% CI: 3.001-57.34), pricked+uv eggs (EMM: 50.026%; 44.37-fold; 95% CI: 10.67-184.4), and UV only eggs (EMM: 41.468%%; 62.69-fold; 17.15-229.1). Natural spawns had the highest percentage of embryos developing to blastula, gastrula, neurula, and tadpole compared to pricked only eggs, pricked+UV eggs, and UV only eggs (See Supplementary Tables 2 and 3).

**Table 3:**
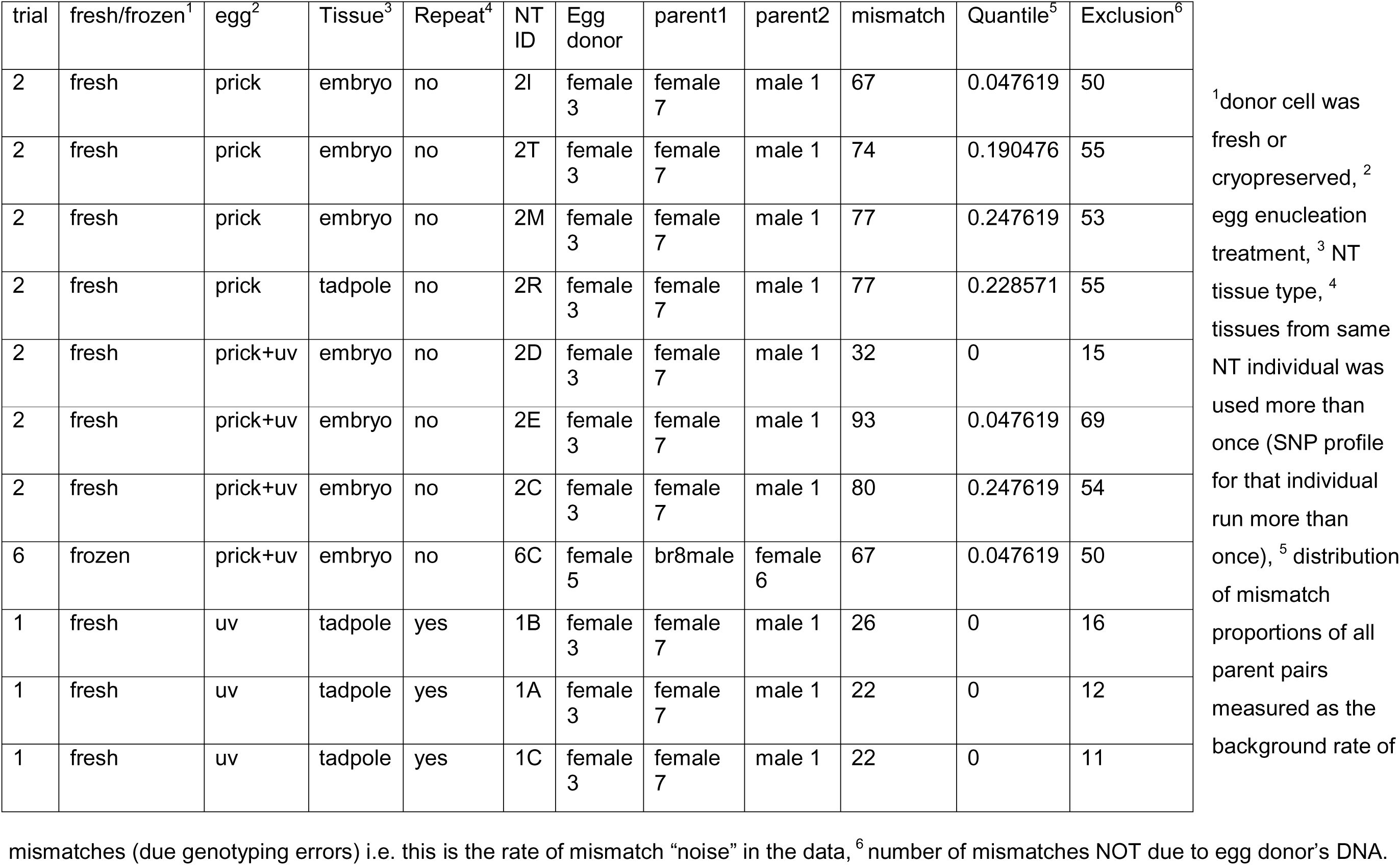
Mismatch and exclusion values counts for the nuclear transplants (NT), parents, and egg donors.

NT embryos from pricked eggs had a significantly higher cleavage rate (See Figure 5), with 3.382-fold and 4.778-fold increases in the percentage of embryos cleaving (EMM: 77.197%) compared to NT embryos from pricked+UV (EMM: 50.266%, 95% CI: 1.016-11.26) and UV only eggs (EMM: 41.468%, 1.674-13.64), respectively. For the percentage of embryos reaching blastula, NT embryos from pricked eggs (EMM: 13.263%) had a significant 5.793-fold and 7.166-fold increase compared to NT embryos from pricked+UV (EMM: 2.572%, 95% CI: 1.074-31.25) and UV only eggs (EMM: 2.089%, 1.909-26.9), respectively. There was no significant difference in the percentage between prick+UV and UV eggs for cleavage (2.684-fold; 95% CI: 6.159-116.9) and blastula (1.237-fold; 95% CI: 0.2446-6.257). For gastrula, there was no significant difference in the percentage of nuclear transplants between UV and prick+UV eggs (1.363-fold; 0.1303-14.25). No NT embryos from pricked only eggs developed beyond blastula.

There was a significant reduction in the percentage of NT embryos that progressed from cleavage to blastula across all egg treatments (See Figure 5) with OR and 95% CI as follows: pricked eggs (OR: 22.14; 95% CI 5.729-85.55), pricked+UV eggs (OR: 37.92; 95% CI: 7.786-184.7), and UV only eggs (OR: 33.20; 95% CI: 11.7-94.2). However, there was no significant decrease in the percentage of NT embryos that progressed from blastula to gastrula for Prick+UV (OR: 2.991; 95% CI: 0.246-36.43) eggs and UV eggs (OR: 1.774; 95% CI: 0.45-6.994) in NT embryos derived from cryopreserved cells. There was no significant drop in the percentage of surviving embryos between gastrula and neurula for UV treated eggs (OR: 4.883; 95% CI: 0.488-48.82).

### SNP Analysis

Nuclear transplants had mismatches between 22-930 (Table 3). The average genotype error rate amongst all nuclear transplants was 0.026384 (Table 4). In error rate (mismatch) simulations, 2.5% of the simulations had mismatch values less than 39, 50% had mismatch values lower than 52, and 97.5% had mismatch values less than 65. Most nuclear transplants had mismatches between 39-66. The 2.5% quantile was accepted as the lower bound of mismatches across the simulations, which suggested that nuclear transplants with less than 39 mismatches are true clones (See Figure 8). This was the boundary accepted as the threshold for determining whether the mismatches of individual NTs indicated that the nuclear genomes of NTs were derived from the injected donor cells (true clones). Mismatches less than 39 were accepted as indicating NT nuclei were true clones of the donor nuclei.

**Figure 8:**
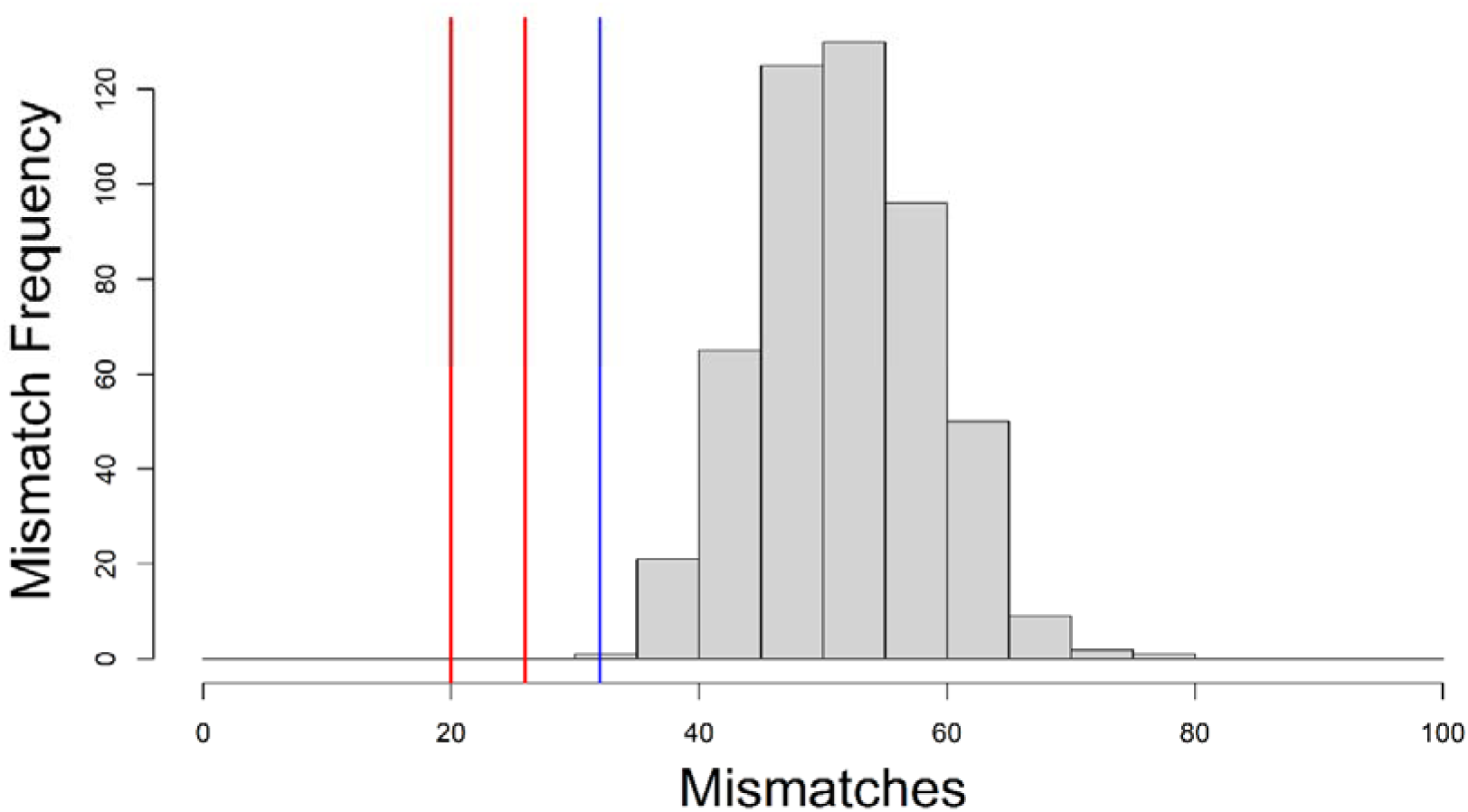
Distribution of 500 mismatch simulations from the dataset of 1108 *L. peronii* SNPs. Red lines are the mismatch counts for the nuclear transplant tadpole from trial 1 (20, 20, 26 mismatches; n = 3 SNP profiles available) and the blue lines are the mismatch count from trial 2 (32 mismatches; n = 1 embryo SNP profile available). The cutoff value was the 2.5% quantile (39 mismatches); values below 39 mismatches taken to indicate genomic identity between donor and NT genome (derived from same genome); values above 39 indicate number of mismatches exceed those predicted by genotyping error.

**Table 4:** Average genotyping Error rates calculated amongst nuclear transplants for each trial and average Error rate across all trials.

| Trial | error rate for each trial | average error rate across all trials |
| --- | --- | --- |
| 4 | 0.03104 | 0.026384 |
| 2 | 0.02592 |  |
| 3 | 0.01909 |  |
| 1 | 0.00137 |  |
| 7 | 0.0545 |  |

From the calculated mismatches, there were two nuclear transplants with low mismatches (see Table 3); these are the nuclear transplant tadpole (from UV only eggs) from trial 1 (26, 20, and 20 for each of three tissue replicates sequenced from this replicate) and a nuclear transplant embryo (from pricked + UV eggs) from trial 2 (32 mismatches between the nuclear transplant and its corresponding donor parent pair). The quantile values were 0 for both the nuclear transplant tadpole and the nuclear transplant embryo (see table 3), which indicated that these two nuclear transplants are true clones. The observed donor parent pairs had the fewest mismatches for their nuclear transplants. Mismatches in the other nuclear transplants from the other trials ranged from 67-93 (for all egg enucleation treatments) and had non-zero quantiles. The non-zero quantiles indicated that those nuclear transplants are not true clones.

Exclusion values are the leftover mismatches after the egg donor’s genotype was taken into account. For nuclear transplant embryos had high exclusion values, those mismatches were not due to the egg donor’s genotype (see Table 3). The higher exclusion values indicated that the egg donor’s DNA likely explains the few of the mismatches. This indicates that these nuclear transplants contain egg donor DNA and are not true clones. The second tadpole from trial 2 had 55/77 mismatches that were not due to egg donor DNA interference, and 22/77 mismatches that are best explained as resulting from the egg donor’s DNA being incorporated into the genome for that tadpole (this tadpole failed much earlier than the tadpole that was a true clone).

## DISCUSSION

This study reported attempts to generate NT embryos using fresh and cryopreserved embryonic cells in an Australian frog. While survival of NTs was low, the development of one embryo to a late-stage tadpole demonstrates proof-of-concept that fresh embryonic cells can support normal development in *L. peronii* up until at least the late tadpole stage. Development of NT embryos from cryopreserved and fresh embryonic cells were significantly lower compared to the natural spawn controls, indicating that egg quality at the time of nuclear transfer was not the cause of low rates of development through to the larval stage. The proportion of surviving embryos decreased for NT embryos derived from fresh and cryopreserved cells for all three egg enucleation treatments. Nevertheless, SNP analysis confirmed that two true clones where produced, indicating that in those replicates the egg’s DNA was successfully enucleated, or did not participate in the development of the nuclear transplants.

Results for the current study can be benchmarked against other reports from the relatively long history of amphibian cloning in non-Australian species. This study reported low development rates of *L. peronii* NT embryos produced from fresh, early embryonic cells (only a single tadpole). Other studies on *X. laevis*, *R. pipiens*, and *A. mexicanum* reported the production of sexually mature adults from fresh embryonic cells (Brothers, 1976; Gurdon et al., 1958; McKinnell, 1962).

The proportion of injected eggs that cleaved for *L. peronii* was 62.73%; this is similar to cleavage rates for *X. laevis* (76%), *R. pipiens* (72%), and *A. mexicanum* (73%) (Briggs et al., 1964; Gurdon, 1962; Hennen, 1970). However, for *R. pipiens*, *A. mexicanum* and *X. laevis*, a higher proportion of injected eggs reached blastula (62% and 96% for A. mexicanum, 72.2%, 72%, 90% for *R. pipiens*, and 62% *for X. laevis*) (Brothers, 1976; Gurdon, 1962; Hennen, 1970; McKinnell, 1962; Signoret et al., 1962; Smith, 1965). Higher transition rates from first cleavage to blastula in these species contrasts with *L. peronii*, in which there was a significant drop in the proportion of embryos surviving between cleavage and blastula (from 67.73% of injected eggs cleaving to 16.9% of attaining blastula). The reasons for variation in development rates after NT between species and studies are difficult to determine. Differences could be due to different methodologies, species specific differences, media types for holding eggs and cells, treatment of the eggs, and cell cycle stage of the donor cell. Two of the major differences among the NT protocols in other species include the means of activation and enucleation of recipient eggs. In *X. laevis*, both activation and enucleation were carried out through UV exposure, and then the donor cell was injected immediately after UV exposure into the egg (Elsdale et al., 1960; Gurdon, 1991). In *R. pipiens* and the *A. mexicanum*, activation and enucleation were performed in two different steps, and then the donor cell nucleus was injected (Briggs & King, 1953; McKinnell, 1978; Signoret et al., 1962). Reasons for these different approaches were in part due to species differences in accessibility of the metaphase pronucleus (egg DNA) during enucleation. Thick jelly layers made access to the pronucleus and physical enucleation challenging in the case of *X. laevis* (Gurdon, 2017), so UV inactivation of the pronucleus became the established, preferred methodology.

In the current study, eggs subjected one of the three egg enucleation treatments (UV, Prick, Prick+UV) were able to support various stages of development, and development to the neurula, and tadpole stage was attained for both UV and Prick treatments (using fresh embryonic cells). However, due to limitations on the number of replicates and small sample size for Prick and Prick+UV treatments, it is uncertain (statistically) which of the three egg treatments were superior for NT with fresh embryonic cells. Another difference between the current study and some of the past studies was that the current study dissociated the entire embryo, without first separating the animal pole, to obtain donor cells. This means that in the current study, cells from both the animal and vegetal poles were injected into the recipient eggs. Many past studies dissociated and used cells from the animal cap in NT (McKinnell, 1978; Signoret et al., 1962). Nevertheless, in one *X. laevis* NT report, there was no difference in the proportion of tadpoles produced from animal pole cells and vegetal cells (Gurdon, 1960).

This study is one of only two to report attempts at nuclear transplantation in amphibians with cryopreserved cells. The other was performed on *Rana temporaria* and *Bufo bufo* using germ cells cryopreserved with DMSO and sucrose. The study reported development of NT embryos to the blastula stage; however, it was not reported if those blastulas progressed, or if the cryoprotectant was removed from the cells post-thaw (Uteshev et al., 2002).

It was not determined from the current study why development after NT using cryopreserved cells from *L. peronii* embryos did not progress as far as with freshly collected embryonic cells. No tadpoles were produced from cryopreserved cells among the three egg enucleation treatments in the current study. Failure to remove cryoprotectant (where DMSO is the cryoprotectant) can affect the donor cells and eggs due to cytotoxicity of DMSO (Adler et al., 2006; Arav et al., 1993). There are few other studies examining the cryopreservation of amphibian early embryonic cells (Lawson et al., 2013; Uteshev et al., 2002), although DMSO proved highly protective of embryonic cells in the current study. In fish, most work with cryopreserved embryonic cells involved the production of chimeras, but did demonstrate that viable embryonic cells were recovered post-thaw because of their incorporation into the germ lines of offspring (Kusuda et al., 2004; Nakagawa et al., 2002).

Studies of cytotoxicity on mammalian and fish embryos revealed that exposure to DMSO resulted in lower survival rates (Kang et al., 2017; Zhang et al., 2005). Even though the cryoprotectant was diluted to a lower concentration before NT in this study, it is possible that the low DMSO concentration was still toxic to the donor cell and egg. However, other aspects of cryoinjury may also be at work. Cryopreservation can comprise DNA integrity, cell membrane, chromatin structure, alter epigenetics, and other cellular structures (Chatterjee et al., 2017; Diwan et al., 2010; Kopeika et al., 2005), which could affect the cell nucleus’s ability to promote development when used in NT.

The use of SNPs in the current study to confirm the generation of true clones has greatly strengthened the outcome of the study and provided confidence in the results. It confirmed the successful generation of true NT clones, but also provided data that suggests that NT may fail in some cases because of incorporation or retention of egg’s DNA in the developing embryo (possibly involving hybridisation of egg and donor cell genomes/chromosomes to some extent. Identifying this as a problem suggests a path forward in future research ((see below) i.e. the improvement of techniques for inactivation and removal of egg donor DNA)). There are a number of possible enucleation and chromosomal aberrations during the NT process that may explain some of the reasons for failure of developing embryos after NT (Figure 9).

**Figure 9.**
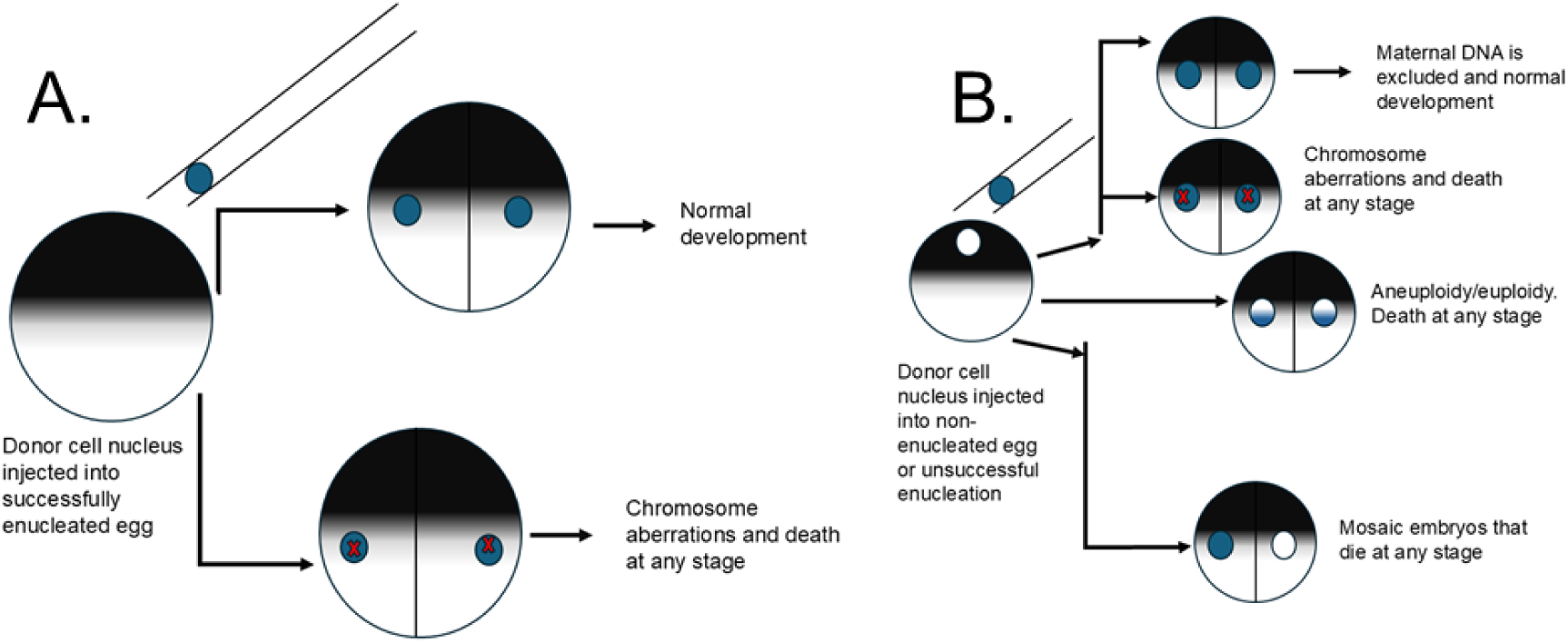
Possible outcomes of NT. 9A has two possible outcomes if the egg was successfully enucleated and the donor cell nucleus (blue) was injected; normal development or chromosome aberrations (red crosses). 9B has three possible outcomes if the egg was not enucleated or if there was unsuccessful (incomplete or partial) enucleation. First, development due to the injected donor cell nucleus can occur and the maternal genome is excluded (white), which can result in normal development, or chromosome aberrations (red crosses). Second, the maternal genome can fuse with the injected donor cell nucleus (blue and white), which can result in aneuploidy/euploidy. Third, mosaic embryos can develop, which results in some cells having the donor cell genome and some cells having the maternal genome.

Using SNP analysis, two true clones were identified through low mismatches between the clones and their corresponding donor parent pairs. One clone was from the prick + UV eggs and one clone was from UV only egg treatments. The low frequency of embryos with low mismatch ratios strongly suggests donor egg DNA was present in many of the embryos with high mismatch levels (since egg donor DNA is a possible source of mismatch DNA where mismatch occurrences are at a sufficiently high frequency to exclude the donor cells as the only source of DNA in the NT embryo genomes). Thus, validating and optimising enucleation is an important question in improving the results of the current study.

In *R. pipiens*, successful enucleation was confirmed by formation of an exovate containing the maternal nucleus as enucleation in ranids was done manually by flicking the maternal nucleus out of the egg with glass needles (Briggs & King, 1957; McKinnell, 1962). However, in *X. laevis* and *A. mexicanum*, eggs were enucleated with UV irradiation and donor cells carrying various mutations were injected into eggs of wild-type phenotype females to allow confirmation genetically that enucleation was successful, and the developing embryo was derived from the injected donor cell (Elsdale et al., 1960; Signoret et al., 1962). UV doses/levels have varied between studies where UV light was used in enucleation. In *X. laevis*, the eggs received either one or two different doses of UV (Elsdale et al., 1960; Gurdon, 1991) and in *A. mexicanum* eggs were exposed to UV for four minutes (Signoret et al., 1962). For *X. laevis*, *R. pipiens*, and *A. mexicanum* NT, successful enucleation of the egg DNA was highly efficient using either physical or UV enucleation, with the result being that the maternal DNA did not participate in the development of nuclear transplant embryos (J. B. Gurdon, 1960; McKinnell, 1978; Signoret et al., 1962). In the current study, it is likely that the *L. peronii* eggs were not exposed long enough to the UV irradiation, resulting in partial or incomplete enucleation.

As stated previously, the evidence of donor egg DNA in the NT embryos suggested that the egg donor’s DNA participated in the development of the NT embryo (with enormous implications for gene expression and a high likelihood of abnormal developmental outcomes). There are a number of possible mechanisms by which this could occur. It is possible that the maternal genome fused with the donor cell nucleus, causing aneuploid/euploid NT embryos or mosaic NT embryos that resulted in combinations of haploid, diploid and even triploid cells (See Figure 9). Studies on fish NT have reported mosaic and/or aneuploid/euploid NT embryos (Bubenshchikova et al., 2005; Depincé et al., 2021; Ju et al., 2003; Kaftanovskaya et al., 2007). It is likely that retaining the maternal DNA would have resulted in developmental abnormalities (Wakamatsu, 2008). In the current study, if the egg DNA fused with the donor cell DNA in hybrid nuclei or even if both genomes were transmitted in the developing embryo, it is likely that development of *L. peronii* NT embryos would fail.

Egg donor DNA could not explain all the mismatches in the NT embryos. An additional possibility that chromosome aberrations (in the donor cell genome) resulted in genetic changes that changed the SNPs of the NT embryos (through deletions, mutations or inversions). In *R. pipiens* NT, chromosome aberrations have been reported, which could have resulted in chromosome deletions, additions, and translocations (Briggs et al., 1964; DiBerardino, 1979; King & Briggs, 1956). If chromosome aberrations occurred in *L. peronii* nuclear transplants, this would have caused mismatches in SNPs between the NT embryo and the donor cell DNA. Although SNP data has proved to be a valuable tool for identifying the genetic origin of nuclear DNA in NTs in the current study, it is likely that SNPs may not be enough to confirm if the NT embryos are true clones if the chromosome aberrations cause mismatches as an artifact of the SNP analysis. In addition to SNPs, karyotyping NT embryos could give additional insight into the chromosome damage that could be occurring in the NT embryos.

Future directions for Australian amphibian cloning for conservation should focus on cell cryopreservation optimization, including investigating DNA damage and epigenetic alterations), investigating the effect of cell cycle stages of the injected donor cells on development as well as maturation/mitosis/meiosis promoting factor (MPF) activity in recipient eggs at the time of NT and determining nuclei damage during transplantation, and optimizing enucleation and/or skipping enucleation the enucleation step altogether.

Since no *L. peronii* NT embryos derived from cryopreserved cells made it to tadpole stage, it is likely there could have been DNA damage and/or epigenetic alterations in cryopreserved embryonic cells. In fish cell cryopreservation studies, DNA strand breaks and epigenetic alterations were observed in primordial germ cells post-thaw (Riesco et al., 2012; Riesco & Robles, 2013). Future work on amphibian embryonic cell cryopreservation should investigate DNA damage and epigenetic alterations in post-thaw cells, and test different additives to the cryoprotectant medium to improve post thaw viability.

Future directions should consider cell cycle phases during NT and the role of maturation promoting factor (MPF) in NT outcomes. In NT, the donor cells should be in G1/G0 when injected into enucleated oocytes at the time of activation (Campbell et al., 1996). Alternatively, donor cells that are at G1, G2, or S phase can be injected into activated and enucleated oocytes after the disappearance of the (MPF) activity (Campbell et al., 1996; Wilmut et al., 2002). MPF activity is responsible for the donor cell nucleus membrane breakdown and premature chromosome condensation. When diploid nuclei in G1/G0 are injected into oocytes when MPF activity is high, the DNA condenses. When nuclei that are replicating (S-phase) are injected into oocytes with high MPF activity, the DNA takes on a pulverized like appearance, which results in DNA damage (Campbell, 1999). Mammalian NT that injected G1 cells into oocytes with high MPF activity or injected G1/S/G2 cells into oocytes with decreased MPF activity resulted in an increase of cloned embryos to the blastocyst stage (Campbell et al., 1996).

In amphibians, MPF activity decreases after fertilization, but then increases around the time of cleavage (Sive, 2023). Cell cycle stages in amphibian early embryos such as *Xenopus* blastulas mostly consist of the S, M, and G2 stages (Siefert et al., 2015). In most amphibian NT studies, early embryonic nuclei such as blastula nuclei were used as donors (Elsdale et al., 1960; McKinnell, 1962; Signoret et al., 1962), so the donor cell nuclei were likely to be in either S, M, or G2. Cell cycle stage of the donor cell was not investigated in *X. laevis*, *R. pipiens*, and *A. mexicanum* nuclear transplantation. Only one study (on *Bombina orientalis*) (Ellinger, 1978) investigated cell cycle of the donor cell and its effect on NT outcome. Donor cells from all cell cycle stages were able to develop normally; however, MPF activity was not addressed, but the eggs received a donor nucleus no later than 30-35 minutes after activation (Ellinger, 1978). Future studies on amphibian NT should inject eggs (with early embryonic cells) right after activation/enucleation or delay injection to investigate if the yield of NT transplants is affected by such factors as in mammalian NT.

It is noted that no specific nuclear media exist for processing amphibian donor cell nuclei. Since the cell membrane of the donor cell must be broken during NT, this exposes the cell nucleus to the electrolyte solution of the media and can result in damage such as swollen nuclei. The ideal electrolyte solutions that worked for NT in amphibians (*X. laevis*, *R. pipiens*, and *A. mexicanum*) were Modified Barth’s Saline, Niu-Twitty, and Steinberg’s Solution (Elsdale et al., 1960; McKinnell, 1978; Signoret et al., 1962). Slight modifications have been made to these such as addition of spermine, BSA, or increasing the magnesium concentration; however, no research has been done to create a specific nuclear transplantation medium or investigated the effects of physiological media on amphibian cell nuclei for NT (Briggs & King, 1953; Hennen, 1970; McKinnell, 1978; Muggleton-Harris, 1971). In models such as zebrafish and mice, media for holding donor cells have been established for NT (Gao et al., 2003; Huang et al., 2003; Lee et al., 2002). Future work should investigate establishing a nuclear medium for amphibian donor cells to be used in NT.

For optimizing enucleation, optimising the UV dose during enucleation could be an important aspect of optimising the NT protocols. Simply increasing the length of time of UV exposure could improve enucleation. However, there is an alternative approach that could investigated: this would be to skip enucleation entirely during the nuclear transfer. In fish NT, there have been reports of true clones being produced with non-enucleated eggs and diploidised eggs (Bubenshchikova et al., 2005; Depincé et al., 2021; Kaftanovskaya et al., 2007). In non-enucleated and diploidised eggs used as recipients for NT, there are instances where the maternal genome did not participate in the development of the NT embryos. The rationale for skipping enucleation in fish NT is due to the presence of yolk reserves and the chorion, which can make enucleation difficult in fish eggs (Depincé et al., 2021). Despite the eggs retaining the maternal DNA, fertile clones that only had donor cell DNA were produced. However, the mechanism of how the maternal nucleus did not participate in NT is unknown (Bubenshchikova et al., 2005, 2007). Future work in amphibian NT should consider using non-enucleated and diploidised eggs to improve success in NT.

## CONCLUSION

The results of the current study confirmed the production of larvae (hind-limb tadpole) by NT in the temperate Australian frog, *Limnodynastes peronii*. Although it was demonstrated that fresh embryonic cells of *L. peronii* can support development of NT embryos to the late tadpole stage, no tadpoles were produced from cryopreserved embryonic cells. Cell nuclei injected into pricked only eggs had higher cleavage rates than other treatments, but enucleation was not successful in most nuclear transplants. Improvement of enucleation procedures for *L. peronii* donor eggs is likely to lead to higher development rates and ultimately to the production of clones that complete development through metamorphosis to the adult phase. Overall, further optimization is needed to create a consistent cloning model that can be applied to amphibian conservation.

