## Supplementary files for "Developing a nuclear transplantation model for Australian amphibian conservation using a common species of Australian Frog (*Limnodynastes peronii*)"

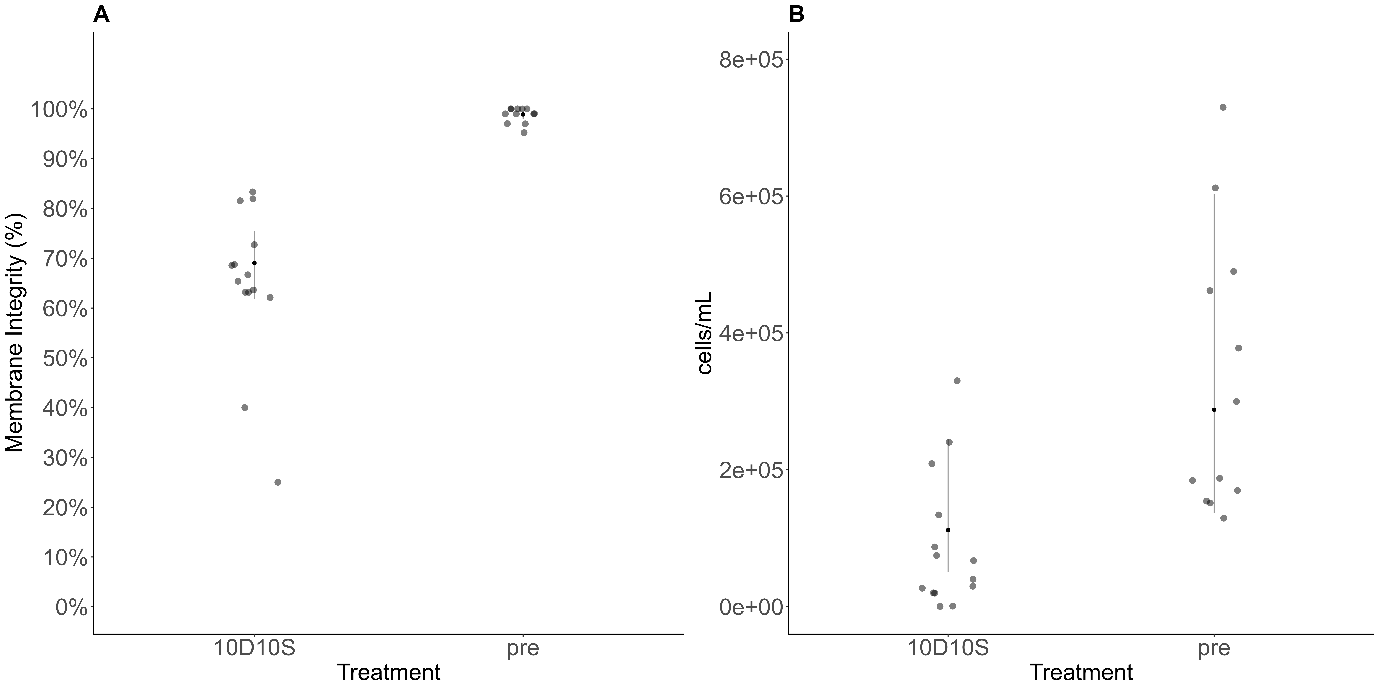

Figure 1: Membrane integrity (1A) and cell concentration (cells/mL) (1B) of cryopreserved (10D10S) and non-cryopreserved (pre-freeze) donor cells used in NT. 1A generated from a GLMM binomial. Black dots are EMMs and black squares are the raw data 2B produce from a zero inflated negative binomial GLMM (black dots are EMMs and grey dots raw data).

Supplementary Table 1: Odds ratios and 95% confidence intervals for development of *L. peronii* nuclear transplants (NTs) from fresh embryonic cells.

| NT Development from Fresh Embryonic Cells | | | | | | | |
| --- | --- | --- | --- | --- | --- | --- | --- |
| reverse contrast | odds ratio | LCL^1^ | UCL^2^ | contrast | odds ratio | LCL^1^ | UCL^2^ |
| NT_UV cleavage / natural_spawn cleavage | 0.05* | 0.007 | 0.334 | natural_spawn cleavage / NT_UV cleavage | 20.141* | 2.991 | 135.651 |
| natural_spawn blastula / natural_spawn cleavage | 1.002 | 0.11 | 9.106 | natural_spawn cleavage / natural_spawn blastula | 0.998 | 0.11 | 9.062 |
| natural_spawn blastula / NT_UV cleavage | 20.189* | 2.995 | 136.086 | natural_spawn cleavage / NT_UV blastula | 166.802* | 23.133 | 1202.713 |
| NT_UV blastula / natural_spawn cleavage | 0.006* | 0.001 | 0.043 | natural_spawn cleavage / natural_spawn gastrula | 1.002 | 0.11 | 9.092 |
| NT_UV blastula / NT_UV cleavage | 0.121* | 0.026 | 0.551 | natural_spawn cleavage / NT_UV gastrula | 341.688* | 44.891 | 2600.764 |
| NT_UV blastula / natural_spawn blastula | 0.006* | 0.001 | 0.043 | natural_spawn cleavage / natural_spawn neurula | 1 | 0.11 | 9.082 |
| natural_spawn gastrula / natural_spawn cleavage | 0.998 | 0.11 | 9.062 | natural_spawn cleavage / NT_UV neurula | 505.192* | 64.118 | 3980.483 |
| natural_spawn gastrula / NT_UV cleavage | 20.108* | 2.987 | 135.353 | natural_spawn cleavage / natural_spawn tadpole | 0.997 | 0.11 | 9.055 |
| natural_spawn gastrula / natural_spawn blastula | 0.996 | 0.11 | 9.045 | natural_spawn cleavage / NT_UV tadpole | 4001.411* | 236.847 | 67601.96 |
| natural_spawn gastrula / NT_UV blastula | 166.525* | 23.109 | 1200.004 | NT_UV cleavage / natural_spawn blastula | 0.05* | 0.007 | 0.334 |
| NT_UV gastrula / natural_spawn cleavage | 0.003* | 0 | 0.022 | NT_UV cleavage / NT_UV blastula | 8.282* | 1.816 | 37.757 |
| NT_UV gastrula / NT_UV cleavage | 0.059* | 0.012 | 0.288 | NT_UV cleavage / natural_spawn gastrula | 0.05* | 0.007 | 0.335 |
| NT_UV gastrula / natural_spawn blastula | 0.003* | 0 | 0.022 | NT_UV cleavage / NT_UV gastrula | 16.965* | 3.478 | 82.745 |
| NT_UV gastrula / NT_UV blastula | 0.488 | 0.096 | 2.489 | NT_UV cleavage / natural_spawn neurula | 0.05* | 0.007 | 0.334 |
| NT_UV gastrula / natural_spawn gastrula | 0.003* | 0 | 0.022 | NT_UV cleavage / NT_UV neurula | 25.083* | 4.87 | 129.188 |
| natural_spawn neurula / natural_spawn cleavage | 1 | 0.11 | 9.077 | NT_UV cleavage / natural_spawn tadpole | 0.049* | 0.007 | 0.334 |
| natural_spawn neurula / NT_UV cleavage | 20.136* | 2.99 | 135.608 | NT_UV cleavage / NT_UV tadpole | 198.668* | 16.402 | 2406.387 |
| natural_spawn neurula / natural_spawn blastula | 0.997 | 0.11 | 9.06 | natural_spawn blastula / NT_UV blastula | 167.2* | 23.168 | 1206.642 |
| natural_spawn neurula / NT_UV blastula | 166.76* | 23.13 | 1202.312 | natural_spawn blastula / natural_spawn gastrula | 1.004 | 0.111 | 9.118 |
| natural_spawn neurula / natural_spawn gastrula | 1.001 | 0.11 | 9.09 | natural_spawn blastula / NT_UV gastrula | 342.505* | 44.96 | 2609.22 |
| natural_spawn neurula / NT_UV gastrula | 341.603* | 44.883 | 2599.901 | natural_spawn blastula / natural_spawn neurula | 1.003 | 0.11 | 9.108 |
| NT_UV neurula / natural_spawn cleavage | 0.002* | 0 | 0.016 | natural_spawn blastula / NT_UV neurula | 506.399* | 64.219 | 3993.235 |
| NT_UV neurula / NT_UV cleavage | 0.04* | 0.008 | 0.205 | natural_spawn blastula / natural_spawn tadpole | 0.999 | 0.11 | 9.081 |
| NT_UV neurula / natural_spawn blastula | 0.002* | 0 | 0.016 | natural_spawn blastula / NT_UV tadpole | 4010.972* | 237.238 | 67813.21 |
| NT_UV neurula / NT_UV blastula | 0.33 | 0.061 | 1.781 | NT_UV blastula / natural_spawn gastrula | 0.006* | 0.001 | 0.043 |
| NT_UV neurula / natural_spawn gastrula | 0.002* | 0 | 0.016 | NT_UV blastula / NT_UV gastrula | 2.048 | 0.402 | 10.443 |
| NT_UV neurula / NT_UV gastrula | 0.676 | 0.118 | 3.875 | NT_UV blastula / natural_spawn neurula | 0.006* | 0.001 | 0.043 |
| NT_UV neurula / natural_spawn neurula | 0.002* | 0 | 0.016 | NT_UV blastula / NT_UV neurula | 3.029 | 0.562 | 16.335 |
| natural_spawn tadpole / natural_spawn cleavage | 1.003 | 0.11 | 9.116 | NT_UV blastula / natural_spawn tadpole | 0.006* | 0.001 | 0.043 |
| natural_spawn tadpole / NT_UV cleavage | 20.208* | 2.997 | 136.254 | NT_UV blastula / NT_UV tadpole | 23.989* | 1.945 | 295.827 |
| natural_spawn tadpole / natural_spawn blastula | 1.001 | 0.11 | 9.098 | natural_spawn gastrula / NT_UV gastrula | 341.122* | 44.843 | 2594.93 |
| natural_spawn tadpole / NT_UV blastula | 167.354* | 23.182 | 1208.161 | natural_spawn gastrula / natural_spawn neurula | 0.999 | 0.11 | 9.064 |
| natural_spawn tadpole / natural_spawn gastrula | 1.005 | 0.111 | 9.128 | natural_spawn gastrula / NT_UV neurula | 504.355* | 64.047 | 3971.691 |
| natural_spawn tadpole / NT_UV gastrula | 342.82* | 44.986 | 2612.487 | natural_spawn gastrula / natural_spawn tadpole | 0.995 | 0.11 | 9.038 |
| natural_spawn tadpole / natural_spawn neurula | 1.004 | 0.11 | 9.118 | natural_spawn gastrula / NT_UV tadpole | 3994.779* | 236.573 | 67456.04 |
| natural_spawn tadpole / NT_UV neurula | 506.866* | 64.258 | 3998.159 | NT_UV gastrula / natural_spawn neurula | 0.003* | 0 | 0.022 |
| NT_UV tadpole / natural_spawn cleavage | 0* | 0 | 0.004 | NT_UV gastrula / NT_UV neurula | 1.479 | 0.258 | 8.472 |
| NT_UV tadpole / NT_UV cleavage | 0.005* | 0 | 0.061 | NT_UV gastrula / natural_spawn tadpole | 0.003* | 0 | 0.022 |
| NT_UV tadpole / natural_spawn blastula | 0* | 0 | 0.004 | NT_UV gastrula / NT_UV tadpole | 11.711 | 0.912 | 150.303 |
| NT_UV tadpole / NT_UV blastula | 0.042* | 0.003 | 0.514 | natural_spawn neurula / NT_UV neurula | 505.066* | 64.107 | 3979.178 |
| NT_UV tadpole / natural_spawn gastrula | 0* | 0 | 0.004 | natural_spawn neurula / natural_spawn tadpole | 0.996 | 0.11 | 9.053 |
| NT_UV tadpole / NT_UV gastrula | 0.085 | 0.007 | 1.096 | natural_spawn neurula / NT_UV tadpole | 4000.41* | 236.804 | 67580.25 |
| NT_UV tadpole / natural_spawn neurula | 0* | 0 | 0.004 | NT_UV neurula / natural_spawn tadpole | 0.002* | 0 | 0.016 |
| NT_UV tadpole / NT_UV neurula | 0.126 | 0.009 | 1.687 | NT_UV neurula / NT_UV tadpole | 7.921 | 0.593 | 105.835 |
| NT_UV tadpole / natural_spawn tadpole | 0* | 0 | 0.004 | natural_spawn tadpole / NT_UV tadpole | 4014.667* | 237.391 | 67894.66 |

^1^Lower confidence limit, ^2^Upper confidence limit, * Significant pairwise comparisons

Supplementary Table 2: Estimated Marginal Means for development of *L. peronii* nuclear transplants from cryopreserved embryonic cells from different egg treatments.

| Egg treatment | stage_reached | EMMEANS (%) |
| --- | --- | --- |
| natural_spawn | cleavage | 97.798 |
| natural_spawn | blastula | 97.798 |
| natural_spawn | gastrula | 97.798 |
| natural_spawn | neurula | 97.798 |
| natural_spawn | tadpole | 95.892 |
| Prick | cleavage | 77.197 |
| Prick | blastula | 13.263 |
| Prick | gastrula | 0.000 |
| Prick | neurula | 0.000 |
| Prick | tadpole | 0.000 |
| Prick_UV | cleavage | 50.026 |
| Prick_UV | blastula | 2.572 |
| Prick_UV | gastrula | 0.875 |
| Prick_UV | neurula | 0.000 |
| Prick_UV | tadpole | 0.000 |
| UV | cleavage | 41.468 |
| UV | blastula | 2.089 |
| UV | gastrula | 1.189 |
| UV | neurula | 0.246 |
| UV | tadpole | 0.000 |

Supplementary Table 3: Odds ratios and 95% confidence intervals for development of *L. peronii* nuclear transplants (NTs) from cryopreserved embryonic cells.

| Development of Nuclear Transplants from Cryopreserved cells | | | | | | | |
| --- | --- | --- | --- | --- | --- | --- | --- |
| reverse contrast | odds ratio | LCL^1^ | UCL^2,3^ | contrast | odds ratio | LCL^1^ | UCL^2,3^ |
| NT_Prick cleavage / natural_spawn cleavage | 0.07623* | 0.01744 | 0.3332 | natural_spawn cleavage / NT_Prick cleavage | 1.312e+01* | 3.001 | 57.34 |
| NT_Prick_UV cleavage / natural_spawn cleavage | 0.02254* | 0.005423 | 0.09368 | natural_spawn cleavage / NT_Prick_UV cleavage | 4.437e+01* | 10.67 | 184.4 |
| NT_Prick_UV cleavage / NT_Prick cleavage | 0.2957* | 0.08883 | 0.9843 | natural_spawn cleavage / NT_UV cleavage | 6.269e+01* | 17.15 | 229.1 |
| NT_UV cleavage / natural_spawn cleavage | 0.01595* | 0.004365 | 0.0583 | natural_spawn cleavage / natural_spawn blastula | 1 | 0.1926 | 5.191 |
| NT_UV cleavage / NT_Prick cleavage | 0.2093* | 0.07332 | 0.5973 | natural_spawn cleavage / NT_Prick blastula | 2.904e+02* | 60.97 | 1384 |
| NT_UV cleavage / NT_Prick_UV cleavage | 0.7077 | 0.2709 | 1.849 | natural_spawn cleavage / NT_Prick_UV blastula | 1.682e+03* | 272.1 | 10400 |
| natural_spawn blastula / natural_spawn cleavage | 1 | 0.1926 | 5.191 | natural_spawn cleavage / NT_UV blastula | 2.081e+03* | 463.2 | 9352 |
| natural_spawn blastula / NT_Prick cleavage | 13.12* | 3.001 | 57.34 | natural_spawn cleavage / natural_spawn gastrula | 1 | 0.1926 | 5.191 |
| natural_spawn blastula / NT_Prick_UV cleavage | 44.37* | 10.67 | 184.4 | natural_spawn cleavage / NT_Prick gastrula | 2.41E+14 | 2.22x10^-16^ | Inf |
| natural_spawn blastula / NT_UV cleavage | 62.69* | 17.15 | 229.1 | natural_spawn cleavage / NT_Prick_UV gastrula | 5.031e+03* | 448.1 | 56500 |
| NT_Prick blastula / natural_spawn cleavage | 0.003443* | 0.000723 | 0.0164 | natural_spawn cleavage / NT_UV gastrula | 3.692e+03* | 735.2 | 18540 |
| NT_Prick blastula / NT_Prick cleavage | 0.04517* | 0.01169 | 0.1746 | natural_spawn cleavage / natural_spawn neurula | 1 | 0.1926 | 5.191 |
| NT_Prick blastula / NT_Prick_UV cleavage | 0.1528* | 0.04266 | 0.547 | natural_spawn cleavage / NT_Prick neurula | 2.07E+13 | 2.22x10^-16^ | Inf |
| NT_Prick blastula / NT_UV cleavage | 0.2158* | 0.06941 | 0.6711 | natural_spawn cleavage / NT_Prick_UV neurula | 9.57E+13 | 2.22x10^-16^ | Inf |
| NT_Prick blastula / natural_spawn blastula | 0.003443* | 0.000723 | 0.0164 | natural_spawn cleavage / NT_UV neurula | 1.803e+04* | 1670 | 194700 |
| NT_Prick_UV blastula / natural_spawn cleavage | 0.0005944* | 9.61E-05 | 0.003675 | natural_spawn cleavage / natural_spawn tadpole | 1.903 | 0.4238 | 8.541 |
| NT_Prick_UV blastula / NT_Prick cleavage | 0.007798* | 0.001504 | 0.04042 | natural_spawn cleavage / NT_Prick tadpole | 3.94E+12 | 2.22x10^-16^ | Inf |
| NT_Prick_UV blastula / NT_Prick_UV cleavage | 0.02637* | 0.005415 | 0.1284 | natural_spawn cleavage / NT_Prick_UV tadpole | 4.22E+13 | 2.22x10^-16^ | Inf |
| NT_Prick_UV blastula / NT_UV cleavage | 0.03726* | 0.008551 | 0.1624 | natural_spawn cleavage / NT_UV tadpole | 1.5E+12 | 2.22x10^-16^ | Inf |
| NT_Prick_UV blastula / natural_spawn blastula | 0.0005944* | 9.61E-05 | 0.003675 | NT_Prick cleavage / NT_Prick_UV cleavage | 3.382e+00* | 1.016 | 11.26 |
| NT_Prick_UV blastula / NT_Prick blastula | 0.1726* | 0.032 | 0.9312 | NT_Prick cleavage / NT_UV cleavage | 4.778e+00* | 1.674 | 13.64 |
| NT_UV blastula / natural_spawn cleavage | 0.0004805* | 0.000107 | 0.002159 | NT_Prick cleavage / natural_spawn blastula | 7.623e-02* | 0.01744 | 0.3332 |
| NT_UV blastula / NT_Prick cleavage | 0.006303* | 0.001754 | 0.02265 | NT_Prick cleavage / NT_Prick blastula | 2.214e+01* | 5.729 | 85.55 |
| NT_UV blastula / NT_Prick_UV cleavage | 0.02132* | 0.006456 | 0.07039 | NT_Prick cleavage / NT_Prick_UV blastula | 1.282e+02* | 24.74 | 664.8 |
| NT_UV blastula / NT_UV cleavage | 0.03012* | 0.01061 | 0.08549 | NT_Prick cleavage / NT_UV blastula | 1.586e+02* | 44.15 | 570.1 |
| NT_UV blastula / natural_spawn blastula | 0.0004805* | 0.000107 | 0.002159 | NT_Prick cleavage / natural_spawn gastrula | 7.623e-02* | 0.01744 | 0.3332 |
| NT_UV blastula / NT_Prick blastula | 0.1395* | 0.03717 | 0.5239 | NT_Prick cleavage / NT_Prick gastrula | 1.84E+13 | 2.22x10^-16^ | Inf |
| NT_UV blastula / NT_Prick_UV blastula | 0.8084 | 0.1598 | 4.088 | NT_Prick cleavage / NT_Prick_UV gastrula | 3.835e+02* | 38.88 | 3783 |
| natural_spawn gastrula / natural_spawn cleavage | 1 | 0.1926 | 5.191 | NT_Prick cleavage / NT_UV gastrula | 2.814e+02* | 68.82 | 1151 |
| natural_spawn gastrula / NT_Prick cleavage | 13.12* | 3.001 | 57.34 | NT_Prick cleavage / natural_spawn neurula | 7.623e-02* | 0.01744 | 0.3332 |
| natural_spawn gastrula / NT_Prick_UV cleavage | 44.37* | 10.67 | 184.4 | NT_Prick cleavage / NT_Prick neurula | 1.58E+12 | 2.22x10^-16^ | Inf |
| natural_spawn gastrula / NT_UV cleavage | 62.69* | 17.15 | 229.1 | NT_Prick cleavage / NT_Prick_UV neurula | 7.29E+12 | 2.22x10^-16^ | Inf |
| natural_spawn gastrula / natural_spawn blastula | 1 | 0.1926 | 5.191 | NT_Prick cleavage / NT_UV neurula | 1.374e+03* | 145.4 | 12990 |
| natural_spawn gastrula / NT_Prick blastula | 290.4* | 60.97 | 1384 | NT_Prick cleavage / natural_spawn tadpole | 1.450e-01* | 0.03909 | 0.5381 |
| natural_spawn gastrula / NT_Prick_UV blastula | 1682* | 272.1 | 10400 | NT_Prick cleavage / NT_Prick tadpole | 3E+11 | 2.22x10^-16^ | Inf |
| natural_spawn gastrula / NT_UV blastula | 2081* | 463.2 | 9352 | NT_Prick cleavage / NT_Prick_UV tadpole | 3.21E+12 | 2.22x10^-16^ | Inf |
| NT_Prick gastrula / natural_spawn cleavage | 4.15E-15 | 2.22x10^-16^ | Inf | NT_Prick cleavage / NT_UV tadpole | 1.14E+11 | 2.22x10^-16^ | Inf |
| NT_Prick gastrula / NT_Prick cleavage | 5.45E-14 | 2.22x10^-16^ | Inf | NT_Prick_UV cleavage / NT_UV cleavage | 1.413 | 0.5409 | 3.691 |
| NT_Prick gastrula / NT_Prick_UV cleavage | 1.84E-13 | 2.22x10^-16^ | Inf | NT_Prick_UV cleavage / natural_spawn blastula | 2.254e-02* | 0.005423 | 0.09368 |
| NT_Prick gastrula / NT_UV cleavage | 2.60E-13 | 2.22x10^-16^ | Inf | NT_Prick_UV cleavage / NT_Prick blastula | 6.546e+00* | 1.828 | 23.44 |
| NT_Prick gastrula / natural_spawn blastula | 4.15E-15 | 2.22x10^-16^ | Inf | NT_Prick_UV cleavage / NT_Prick_UV blastula | 3.792e+01* | 7.786 | 184.7 |
| NT_Prick gastrula / NT_Prick blastula | 1.21E-12 | 2.22x10^-16^ | Inf | NT_Prick_UV cleavage / NT_UV blastula | 4.691e+01* | 14.21 | 154.9 |
| NT_Prick gastrula / NT_Prick_UV blastula | 6.99E-12 | 2.22x10^-16^ | Inf | NT_Prick_UV cleavage / natural_spawn gastrula | 2.254e-02* | 0.005423 | 0.09368 |
| NT_Prick gastrula / NT_UV blastula | 8.65E-12 | 2.22x10^-16^ | Inf | NT_Prick_UV cleavage / NT_Prick gastrula | 5.43E+12 | 2.22E-16 | Inf |
| NT_Prick gastrula / natural_spawn gastrula | 4.15E-15 | 2.22x10^-16^ | Inf | NT_Prick_UV cleavage / NT_Prick_UV gastrula | 1.134e+02* | 12.02 | 1070 |
| NT_Prick_UV gastrula / natural_spawn cleavage | 0.0001988* | 1.77E-05 | 0.002232 | NT_Prick_UV cleavage / NT_UV gastrula | 8.322e+01* | 21.96 | 315.4 |
| NT_Prick_UV gastrula / NT_Prick cleavage | 0.002607* | 0.000264 | 0.02572 | NT_Prick_UV cleavage / natural_spawn neurula | 2.254e-02* | 0.005423 | 0.09368 |
| NT_Prick_UV gastrula / NT_Prick_UV cleavage | 0.008818* | 0.000935 | 0.08321 | NT_Prick_UV cleavage / NT_Prick neurula | 4.66E+11 | 2.22x10^-16^ | Inf |
| NT_Prick_UV gastrula / NT_UV cleavage | 0.01246* | 0.001426 | 0.1088 | NT_Prick_UV cleavage / NT_Prick_UV neurula | 2.16E+12 | 2.22x10^-16^ | Inf |
| NT_Prick_UV gastrula / natural_spawn blastula | 0.0001988* | 1.77E-05 | 0.002232 | NT_Prick_UV cleavage / NT_UV neurula | 4.064e+02* | 45.04 | 3667 |
| NT_Prick_UV gastrula / NT_Prick blastula | 0.05773* | 0.005685 | 0.5861 | NT_Prick_UV cleavage / natural_spawn tadpole | 4.288e-02* | 0.01223 | 0.1503 |
| NT_Prick_UV gastrula / NT_Prick_UV blastula | 0.3344 | 0.02745 | 4.074 | NT_Prick_UV cleavage / NT_Prick tadpole | 8.88E+10 | 2.22x10^-16^ | Inf |
| NT_Prick_UV gastrula / NT_UV blastula | 0.4137 | 0.04267 | 4.01 | NT_Prick_UV cleavage / NT_Prick_UV tadpole | 9.5E+11 | 2.22x10^-16^ | Inf |
| NT_Prick_UV gastrula / natural_spawn gastrula | 0.0001988* | 1.77E-05 | 0.002232 | NT_Prick_UV cleavage / NT_UV tadpole | 3.37E+10 | 2.22x10^-16^ | Inf |
| NT_Prick_UV gastrula / NT_Prick gastrula | 4.79E+10 | 2.22x10^-16^ | Inf | NT_UV cleavage / natural_spawn blastula | 1.595e-02* | 0.004365 | 0.0583 |
| NT_UV gastrula / natural_spawn cleavage | 0.0002708* | 5.39E-05 | 0.00136 | NT_UV cleavage / NT_Prick blastula | 4.633e+00* | 1.49 | 14.41 |
| NT_UV gastrula / NT_Prick cleavage | 0.003553* | 0.000869 | 0.01453 | NT_UV cleavage / NT_Prick_UV blastula | 2.684e+01* | 6.159 | 116.9 |
| NT_UV gastrula / NT_Prick_UV cleavage | 0.01202* | 0.003171 | 0.04554 | NT_UV cleavage / NT_UV blastula | 3.320e+01* | 11.7 | 94.24 |
| NT_UV gastrula / NT_UV cleavage | 0.01698* | 0.005121 | 0.05629 | NT_UV cleavage / natural_spawn gastrula | 1.595e-02* | 0.004365 | 0.0583 |
| NT_UV gastrula / natural_spawn blastula | 0.0002708* | 5.39E-05 | 0.00136 | NT_UV cleavage / NT_Prick gastrula | 3.84E+12 | 2.22x10^-16^ | Inf |
| NT_UV gastrula / NT_Prick blastula | 0.07866* | 0.01847 | 0.3349 | NT_UV cleavage / NT_Prick_UV gastrula | 8.026e+01* | 9.187 | 701.2 |
| NT_UV gastrula / NT_Prick_UV blastula | 0.4557 | 0.08117 | 2.558 | NT_UV cleavage / NT_UV gastrula | 5.890e+01* | 17.77 | 195.3 |
| NT_UV gastrula / NT_UV blastula | 0.5637 | 0.143 | 2.222 | NT_UV cleavage / natural_spawn neurula | 1.595e-02* | 0.004365 | 0.0583 |
| NT_UV gastrula / natural_spawn gastrula | 0.0002708* | 5.39E-05 | 0.00136 | NT_UV cleavage / NT_Prick neurula | 3.3E+11 | 2.22E-16 | Inf |
| NT_UV gastrula / NT_Prick gastrula | 6.52E+10 | 2.22x10^-16^ | Inf | NT_UV cleavage / NT_Prick_UV neurula | 1.53E+12 | 2.22E-16 | Inf |
| NT_UV gastrula / NT_Prick_UV gastrula | 1.363 | 0.1303 | 14.25 | NT_UV cleavage / NT_UV neurula | 2.876e+02* | 34.48 | 2399 |
| natural_spawn neurula / natural_spawn cleavage | 1 | 0.1926 | 5.191 | NT_UV cleavage / natural_spawn tadpole | 3.035e-02* | 0.01004 | 0.09173 |
| natural_spawn neurula / NT_Prick cleavage | 13.12* | 3.001 | 57.34 | NT_UV cleavage / NT_Prick tadpole | 6.29E+10 | 2.22x10^-16^ | Inf |
| natural_spawn neurula / NT_Prick_UV cleavage | 44.37* | 10.67 | 184.4 | NT_UV cleavage / NT_Prick_UV tadpole | 6.73E+11 | 2.22x10^-16^ | Inf |
| natural_spawn neurula / NT_UV cleavage | 62.69* | 17.15 | 229.1 | NT_UV cleavage / NT_UV tadpole | 2.38E+10 | 2.22x10^-16^ | Inf |
| natural_spawn neurula / natural_spawn blastula | 1 | 0.1926 | 5.191 | natural_spawn blastula / NT_Prick blastula | 2.904e+02* | 60.97 | 1384 |
| natural_spawn neurula / NT_Prick blastula | 290.4* | 60.97 | 1384 | natural_spawn blastula / NT_Prick_UV blastula | 1.682e+03* | 272.1 | 10400 |
| natural_spawn neurula / NT_Prick_UV blastula | 1682* | 272.1 | 10400 | natural_spawn blastula / NT_UV blastula | 2.081e+03* | 463.2 | 9352 |
| natural_spawn neurula / NT_UV blastula | 2081* | 463.2 | 9352 | natural_spawn blastula / natural_spawn gastrula | 1 | 0.1926 | 5.191 |
| natural_spawn neurula / natural_spawn gastrula | 1 | 0.1926 | 5.191 | natural_spawn blastula / NT_Prick gastrula | 2.41E+14 | 2.22x10^-16^ | Inf |
| natural_spawn neurula / NT_Prick gastrula | 2.41E+14 | 2.22E-16 | Inf | natural_spawn blastula / NT_Prick_UV gastrula | 5.031e+03* | 448.1 | 56500 |
| natural_spawn neurula / NT_Prick_UV gastrula | 5031* | 448.1 | 56500 | natural_spawn blastula / NT_UV gastrula | 3.692e+03* | 735.2 | 18540 |
| natural_spawn neurula / NT_UV gastrula | 3692* | 735.2 | 18540 | natural_spawn blastula / natural_spawn neurula | 1 | 0.1926 | 5.191 |
| NT_Prick neurula / natural_spawn cleavage | 4.84E-14 | 2.22x10^-16^ | Inf | natural_spawn blastula / NT_Prick neurula | 2.07E+13 | 2.22x10^-16^ | Inf |
| NT_Prick neurula / NT_Prick cleavage | 6.35E-13 | 2.22x10^-16^ | Inf | natural_spawn blastula / NT_Prick_UV neurula | 9.57E+13 | 2.22x10^-16^ | Inf |
| NT_Prick neurula / NT_Prick_UV cleavage | 2.15E-12 | 2.22x10^-16^ | Inf | natural_spawn blastula / NT_UV neurula | 1.803e+04* | 1670 | 194700 |
| NT_Prick neurula / NT_UV cleavage | 3.03E-12 | 2.22x10^-16^ | Inf | natural_spawn blastula / natural_spawn tadpole | 1.903 | 0.4238 | 8.541 |
| NT_Prick neurula / natural_spawn blastula | 4.84E-14 | 2.22x10^-16^ | Inf | natural_spawn blastula / NT_Prick tadpole | 3.94E+12 | 2.22x10^-16^ | Inf |
| NT_Prick neurula / NT_Prick blastula | 1.41E-11 | 2.22x10^-16^ | Inf | natural_spawn blastula / NT_Prick_UV tadpole | 4.22E+13 | 2.22x10^-16^ | Inf |
| NT_Prick neurula / NT_Prick_UV blastula | 8.14E-11 | 2.22x10^-16^ | Inf | natural_spawn blastula / NT_UV tadpole | 1.5E+12 | 2.22x10^-16^ | Inf |
| NT_Prick neurula / NT_UV blastula | 1.01E-10 | 2.22x10^-16^ | Inf | NT_Prick blastula / NT_Prick_UV blastula | 5.793e+00* | 1.074 | 31.25 |
| NT_Prick neurula / natural_spawn gastrula | 4.84E-14 | 2.22x10^-16^ | Inf | NT_Prick blastula / NT_UV blastula | 7.166e+00* | 1.909 | 26.9 |
| NT_Prick neurula / NT_Prick gastrula | 11.65 | 2.22x10^-16^ | Inf | NT_Prick blastula / natural_spawn gastrula | 3.443e-03* | 0.000723 | 0.0164 |
| NT_Prick neurula / NT_Prick_UV gastrula | 2.43E-10 | 2.22x10^-16^ | Inf | NT_Prick blastula / NT_Prick gastrula | 8.29E+11 | 2.22x10^-16^ | Inf |
| NT_Prick neurula / NT_UV gastrula | 1.79E-10 | 2.22x10^-16^ | Inf | NT_Prick blastula / NT_Prick_UV gastrula | 1.732e+01* | 1.706 | 175.9 |
| NT_Prick neurula / natural_spawn neurula | 4.84E-14 | 2.22x10^-16^ | Inf | NT_Prick blastula / NT_UV gastrula | 1.271e+01* | 2.986 | 54.13 |
| NT_Prick_UV neurula / natural_spawn cleavage | 1.05E-14 | 2.22x10^-16^ | Inf | NT_Prick blastula / natural_spawn neurula | 3.443e-03* | 0.000723 | 0.0164 |
| NT_Prick_UV neurula / NT_Prick cleavage | 1.37E-13 | 2.22x10^-16^ | Inf | NT_Prick blastula / NT_Prick neurula | 7.12E+10 | 2.22x10^-16^ | Inf |
| NT_Prick_UV neurula / NT_Prick_UV cleavage | 4.64E-13 | 2.22x10^-16^ | Inf | NT_Prick blastula / NT_Prick_UV neurula | 3.3E+11 | 2.22x10^-16^ | Inf |
| NT_Prick_UV neurula / NT_UV cleavage | 6.55E-13 | 2.22x10^-16^ | Inf | NT_Prick blastula / NT_UV neurula | 6.208e+01* | 6.393 | 602.8 |
| NT_Prick_UV neurula / natural_spawn blastula | 1.05E-14 | 2.22x10^-16^ | Inf | NT_Prick blastula / natural_spawn tadpole | 6.551e-03* | 0.001603 | 0.02677 |
| NT_Prick_UV neurula / NT_Prick blastula | 3.04E-12 | 2.22x10^-16^ | Inf | NT_Prick blastula / NT_Prick tadpole | 1.36E+10 | 2.22x10^-16^ | Inf |
| NT_Prick_UV neurula / NT_Prick_UV blastula | 1.76E-11 | 2.22x10^-16^ | Inf | NT_Prick blastula / NT_Prick_UV tadpole | 1.45E+11 | 2.22x10^-16^ | Inf |
| NT_Prick_UV neurula / NT_UV blastula | 2.18E-11 | 2.22x10^-16^ | Inf | NT_Prick blastula / NT_UV tadpole | 5.15E+09 | 2.22x10^-16^ | Inf |
| NT_Prick_UV neurula / natural_spawn gastrula | 1.05E-14 | 2.22x10^-16^ | Inf | NT_Prick_UV blastula / NT_UV blastula | 1.237 | 0.2446 | 6.257 |
| NT_Prick_UV neurula / NT_Prick gastrula | 2.516 | 2.22x10^-16^ | Inf | NT_Prick_UV blastula / natural_spawn gastrula | 5.944e-04* | 9.61E-05 | 0.003675 |
| NT_Prick_UV neurula / NT_Prick_UV gastrula | 5.26E-11 | 2.22x10^-16^ | Inf | NT_Prick_UV blastula / NT_Prick gastrula | 1.43E+11 | 2.22x10^-16^ | Inf |
| NT_Prick_UV neurula / NT_UV gastrula | 3.86E-11 | 2.22x10^-16^ | Inf | NT_Prick_UV blastula / NT_Prick_UV gastrula | 2.991 | 0.2455 | 36.43 |
| NT_Prick_UV neurula / natural_spawn neurula | 1.05E-14 | 2.22x10^-16^ | Inf | NT_Prick_UV blastula / NT_UV gastrula | 2.195 | 0.3909 | 12.32 |
| NT_Prick_UV neurula / NT_Prick neurula | 0.216 | 2.22E-16 | Inf | NT_Prick_UV blastula / natural_spawn neurula | 5.944e-04* | 9.61E-05 | 0.003675 |
| NT_UV neurula / natural_spawn cleavage | 5.546e-05* | 5.14E-06 | 0.000599 | NT_Prick_UV blastula / NT_Prick neurula | 1.23E+10 | 2.22x10^-16^ | Inf |
| NT_UV neurula / NT_Prick cleavage | 0.0007276* | 7.70E-05 | 0.006877 | NT_Prick_UV blastula / NT_Prick_UV neurula | 5.69E+10 | 2.22x10^-16^ | Inf |
| NT_UV neurula / NT_Prick_UV cleavage | 0.002461* | 0.000273 | 0.0222 | NT_Prick_UV blastula / NT_UV neurula | 10.72 | 0.9168 | 125.3 |
| NT_UV neurula / NT_UV cleavage | 0.003477* | 0.000417 | 0.02901 | NT_Prick_UV blastula / natural_spawn tadpole | 1.131e-03* | 0.000208 | 0.006141 |
| NT_UV neurula / natural_spawn blastula | 5.546e-05* | 5.14E-06 | 0.000599 | NT_Prick_UV blastula / NT_Prick tadpole | 2.34E+09 | 2.22x10^-16^ | Inf |
| NT_UV neurula / NT_Prick blastula | 0.01611* | 0.001659 | 0.1564 | NT_Prick_UV blastula / NT_Prick_UV tadpole | 2.51E+10 | 2.22x10^-16^ | Inf |
| NT_UV neurula / NT_Prick_UV blastula | 0.09331 | 0.007983 | 1.091 | NT_Prick_UV blastula / NT_UV tadpole | 8.88E+08 | 2.22x10^-16^ | Inf |
| NT_UV neurula / NT_UV blastula | 0.1154 | 0.01247 | 1.069 | NT_UV blastula / natural_spawn gastrula | 4.805e-04* | 0.000107 | 0.002159 |
| NT_UV neurula / natural_spawn gastrula | 5.546e-05* | 5.14E-06 | 0.000599 | NT_UV blastula / NT_Prick gastrula | 1.16E+11 | 2.22E-16 | Inf |
| NT_UV neurula / NT_Prick gastrula | 1.34E+10 | 2.22E-16 | Inf | NT_UV blastula / NT_Prick_UV gastrula | 2.417 | 0.2494 | 23.44 |
| NT_UV neurula / NT_Prick_UV gastrula | 0.2791 | 0.01492 | 5.219 | NT_UV blastula / NT_UV gastrula | 1.774 | 0.45 | 6.994 |
| NT_UV neurula / NT_UV gastrula | 0.2048 | 0.02048 | 2.048 | NT_UV blastula / natural_spawn neurula | 4.805e-04* | 0.000107 | 0.002159 |
| NT_UV neurula / natural_spawn neurula | 5.546e-05* | 5.14E-06 | 0.000599 | NT_UV blastula / NT_Prick neurula | 9.93E+09 | 2.22x10^-16^ | Inf |
| NT_UV neurula / NT_Prick neurula | 1.15E+09 | 2.22x10^-16^ | Inf | NT_UV blastula / NT_Prick_UV neurula | 4.6E+10 | 2.22x10^-16^ | Inf |
| NT_UV neurula / NT_Prick_UV neurula | 5.31E+09 | 2.22x10^-16^ | Inf | NT_UV blastula / NT_UV neurula | 8.663 | 0.9359 | 80.19 |
| natural_spawn tadpole / natural_spawn cleavage | 0.5256 | 0.1171 | 2.36 | NT_UV blastula / natural_spawn tadpole | 9.141e-04* | 0.000239 | 0.0035 |
| natural_spawn tadpole / NT_Prick cleavage | 6.895* | 1.859 | 25.58 | NT_UV blastula / NT_Prick tadpole | 1.89E+09 | 2.22x10^-16^ | Inf |
| natural_spawn tadpole / NT_Prick_UV cleavage | 23.32* | 6.653 | 81.74 | NT_UV blastula / NT_Prick_UV tadpole | 2.03E+10 | 2.22x10^-16^ | Inf |
| natural_spawn tadpole / NT_UV cleavage | 32.95* | 10.9 | 99.58 | NT_UV blastula / NT_UV tadpole | 7.18E+08 | 2.22x10^-16^ | Inf |
| natural_spawn tadpole / natural_spawn blastula | 0.5256 | 0.1171 | 2.36 | natural_spawn gastrula / NT_Prick gastrula | 2.41E+14 | 2.22x10^-16^ | Inf |
| natural_spawn tadpole / NT_Prick blastula | 152.7* | 37.36 | 623.7 | natural_spawn gastrula / NT_Prick_UV gastrula | 5.031e+03* | 448.1 | 56500 |
| natural_spawn tadpole / NT_Prick_UV blastula | 884.3* | 162.8 | 4802 | natural_spawn gastrula / NT_UV gastrula | 3.692e+03* | 735.2 | 18540 |
| natural_spawn tadpole / NT_UV blastula | 1094* | 285.7 | 4188 | natural_spawn gastrula / natural_spawn neurula | 1 | 0.1926 | 5.191 |
| natural_spawn tadpole / natural_spawn gastrula | 0.5256 | 0.1171 | 2.36 | natural_spawn gastrula / NT_Prick neurula | 2.07E+13 | 2.22x10^-16^ | Inf |
| natural_spawn tadpole / NT_Prick gastrula | 1.27E+14 | 2.22x10^-16^ | Inf | natural_spawn gastrula / NT_Prick_UV neurula | 9.57E+13 | 2.22x10^-16^ | Inf |
| natural_spawn tadpole / NT_Prick_UV gastrula | 2644* | 259.3 | 26970 | natural_spawn gastrula / NT_UV neurula | 1.803e+04* | 1670 | 194700 |
| natural_spawn tadpole / NT_UV gastrula | 1941* | 448 | 8407 | natural_spawn gastrula / natural_spawn tadpole | 1.903 | 0.4238 | 8.541 |
| natural_spawn tadpole / natural_spawn neurula | 0.5256 | 0.1171 | 2.36 | natural_spawn gastrula / NT_Prick tadpole | 3.94E+12 | 2.22x10^-16^ | Inf |
| natural_spawn tadpole / NT_Prick neurula | 1.09E+13 | 2.22x10^-16^ | Inf | natural_spawn gastrula / NT_Prick_UV tadpole | 4.22E+13 | 2.22x10^-16^ | Inf |
| natural_spawn tadpole / NT_Prick_UV neurula | 5.03x10^13^ | 2.22x10^-16^ | Inf | natural_spawn gastrula / NT_UV tadpole | 1.5E+12 | 2.22x10^-16^ | Inf |
| natural_spawn tadpole / NT_UV neurula | 9476* | 967.7 | 92800 | NT_Prick gastrula / NT_Prick_UV gastrula | 2.09E-11 | 2.22x10^-16^ | Inf |
| NT_Prick tadpole / natural_spawn cleavage | 2.54x10^-13^ | 2.22x10^-16^ | Inf | NT_Prick gastrula / NT_UV gastrula | 1.53E-11 | 2.22x10^-16^ | Inf |
| NT_Prick tadpole / NT_Prick cleavage | 3.33x10^-12^ | 2.22x10^-16^ | Inf | NT_Prick gastrula / natural_spawn neurula | 4.15E-15 | 2.22x10^-16^ | Inf |
| NT_Prick tadpole / NT_Prick_UV cleavage | 1.13x10^-11^ | 2.22x10^-16^ | Inf | NT_Prick gastrula / NT_Prick neurula | 0.08587 | 2.22x10^-16^ | Inf |
| NT_Prick tadpole / NT_UV cleavage | 1.59x10^-11^ | 2.22x10^-16^ | Inf | NT_Prick gastrula / NT_Prick_UV neurula | 0.3975 | 2.22x10^-16^ | Inf |
| NT_Prick tadpole / natural_spawn blastula | 2.54x10^-13^ | 2.22x10^-16^ | Inf | NT_Prick gastrula / NT_UV neurula | 7.49E-11 | 2.22x10^-16^ | Inf |
| NT_Prick tadpole / NT_Prick blastula | 7.37x10^-11^ | 2.22x10^-16^ | Inf | NT_Prick gastrula / natural_spawn tadpole | 7.9E-15 | 2.22x10^-16^ | Inf |
| NT_Prick tadpole / NT_Prick_UV blastula | 4.27x10^-10^ | 2.22x10^-16^ | Inf | NT_Prick gastrula / NT_Prick tadpole | 0.01637 | 2.22x10^-16^ | Inf |
| NT_Prick tadpole / NT_UV blastula | 5.28x10^-10^ | 2.22x10^-16^ | Inf | NT_Prick gastrula / NT_Prick_UV tadpole | 0.1752 | 2.22x10^-16^ | Inf |
| NT_Prick tadpole / natural_spawn gastrula | 2.54x10^-13^ | 2.22x10^-16^ | Inf | NT_Prick gastrula / NT_UV tadpole | 0.006209 | 2.22x10^-16^ | Inf |
| NT_Prick tadpole / NT_Prick gastrula | 61.1 | 2.22x10^-16^ | Inf | NT_Prick_UV gastrula / NT_UV gastrula | 0.7339 | 0.07019 | 7.672 |
| NT_Prick tadpole / NT_Prick_UV gastrula | 1.28x10^-09^ | 2.22x10^-16^ | Inf | NT_Prick_UV gastrula / natural_spawn neurula | 1.988e-04* | 1.77E-05 | 0.002232 |
| NT_Prick tadpole / NT_UV gastrula | 9.37x10^-10^ | 2.22x10^-16^ | Inf | NT_Prick_UV gastrula / NT_Prick neurula | 4.11E+09 | 2.22x10^-16^ | Inf |
| NT_Prick tadpole / natural_spawn neurula | 2.54x10^-13^ | 2.22x10^-16^ | Inf | NT_Prick_UV gastrula / NT_Prick_UV neurula | 1.9E+10 | 2.22x10^-16^ | Inf |
| NT_Prick tadpole / NT_Prick neurula | 5.246 | 2.22x10^-16^ | Inf | NT_Prick_UV gastrula / NT_UV neurula | 3.583 | 0.1916 | 67.02 |
| NT_Prick tadpole / NT_Prick_UV neurula | 24.29 | 2.22x10^-16^ | Inf | NT_Prick_UV gastrula / natural_spawn tadpole | 3.781e-04* | 3.71E-05 | 0.003857 |
| NT_Prick tadpole / NT_UV neurula | 4.58x10^-09^ | 2.22x10^-16^ | Inf | NT_Prick_UV gastrula / NT_Prick tadpole | 7.83E+08 | 2.22x10^-16^ | Inf |
| NT_Prick tadpole / natural_spawn tadpole | 4.83x10^-13^ | 2.22x10^-16^ | Inf | NT_Prick_UV gastrula / NT_Prick_UV tadpole | 8.38E+09 | 2.22x10^-16^ | Inf |
| NT_Prick_UV tadpole / natural_spawn cleavage | 2.37x10^-14^ | 2.22x10^-16^ | Inf | NT_Prick_UV gastrula / NT_UV tadpole | 2.97E+08 | 2.22x10^-16^ | Inf |
| NT_Prick_UV tadpole / NT_Prick cleavage | 3.11x10^-13^ | 2.22x10^-16^ | Inf | NT_UV gastrula / natural_spawn neurula | 2.708e-04* | 5.39E-05 | 0.00136 |
| NT_Prick_UV tadpole / NT_Prick_UV cleavage | 1.05x10^-12^ | 2.22x10^-16^ | Inf | NT_UV gastrula / NT_Prick neurula | 5.6E+09 | 2.22x10^-16^ | Inf |
| NT_Prick_UV tadpole / NT_UV cleavage | 1.49x10^-12^ | 2.22x10^-16^ | Inf | NT_UV gastrula / NT_Prick_UV neurula | 2.59E+10 | 2.22x10^-16^ | Inf |
| NT_Prick_UV tadpole / natural_spawn blastula | 2.37x10^-14^ | 2.22x10^-16^ | Inf | NT_UV gastrula / NT_UV neurula | 4.883 | 0.4884 | 48.82 |
| NT_Prick_UV tadpole / NT_Prick blastula | 6.89x10^-12^ | 2.22x10^-16^ | Inf | NT_UV gastrula / natural_spawn tadpole | 5.153e-04* | 0.000119 | 0.002232 |
| NT_Prick_UV tadpole / NT_Prick_UV blastula | 3.99x10^-11^ | 2.22x10^-16^ | Inf | NT_UV gastrula / NT_Prick tadpole | 1.07E+09 | 2.22x10^-16^ | Inf |
| NT_Prick_UV tadpole / NT_UV blastula | 4.94x10^-11^ | 2.22x10^-16^ | Inf | NT_UV gastrula / NT_Prick_UV tadpole | 1.14E+10 | 2.22x10^-16^ | Inf |
| NT_Prick_UV tadpole / natural_spawn gastrula | 2.37x10^-14^ | 2.22x10^-16^ | Inf | NT_UV gastrula / NT_UV tadpole | 4.05E+08 | 2.22x10^-16^ | Inf |
| NT_Prick_UV tadpole / NT_Prick gastrula | 5.709 | 2.22x10^-16^ | Inf | natural_spawn neurula / NT_Prick neurula | 2.07E+13 | 2.22x10^-16^ | Inf |
| NT_Prick_UV tadpole / NT_Prick_UV gastrula | 1.19x10^-10^ | 2.22x10^-16^ | Inf | natural_spawn neurula / NT_Prick_UV neurula | 9.57E+13 | 2.22x10^-16^ | Inf |
| NT_Prick_UV tadpole / NT_UV gastrula | 8.76x10^-11^ | 2.22x10^-16^ | Inf | natural_spawn neurula / NT_UV neurula | 1.803e+04* | 1670 | 194700 |
| NT_Prick_UV tadpole / natural_spawn neurula | 2.37x10^-14^ | 2.22x10^-16^ | Inf | natural_spawn neurula / natural_spawn tadpole | 1.903 | 0.4238 | 8.541 |
| NT_Prick_UV tadpole / NT_Prick neurula | 0.4902 | 2.22x10^-16^ | Inf | natural_spawn neurula / NT_Prick tadpole | 3.94E+12 | 2.22x10^-16^ | Inf |
| NT_Prick_UV tadpole / NT_Prick_UV neurula | 2.269 | 2.22x10^-16^ | Inf | natural_spawn neurula / NT_Prick_UV tadpole | 4.22E+13 | 2.22x10^-16^ | Inf |
| NT_Prick_UV tadpole / NT_UV neurula | 4.28x10^-10^ | 2.22x10^-16^ | Inf | natural_spawn neurula / NT_UV tadpole | 1.5E+12 | 2.22x10^-16^ | Inf |
| NT_Prick_UV tadpole / natural_spawn tadpole | 4.51x10^-14^ | 2.22x10^-16^ | Inf | NT_Prick neurula / NT_Prick_UV neurula | 4.629 | 2.22x10^-16^ | Inf |
| NT_Prick_UV tadpole / NT_Prick tadpole | 0.09344 | 2.22x10^-16^ | Inf | NT_Prick neurula / NT_UV neurula | 8.72E-10 | 2.22x10^-16^ | Inf |
| NT_UV tadpole / natural_spawn cleavage | 6.69x10^-13^ | 2.22x10^-16^ | Inf | NT_Prick neurula / natural_spawn tadpole | 9.2E-14 | 2.22x10^-16^ | Inf |
| NT_UV tadpole / NT_Prick cleavage | 8.78x10^-12^ | 2.22x10^-16^ | Inf | NT_Prick neurula / NT_Prick tadpole | 0.1906 | 2.22x10^-16^ | Inf |
| NT_UV tadpole / NT_Prick_UV cleavage | 2.97x10^-11^ | 2.22x10^-16^ | Inf | NT_Prick neurula / NT_Prick_UV tadpole | 2.04 | 2.22x10^-16^ | Inf |
| NT_UV tadpole / NT_UV cleavage | 4.19x10^-11^ | 2.22x10^-16^ | Inf | NT_Prick neurula / NT_UV tadpole | 0.07231 | 2.22x10^-16^ | Inf |
| NT_UV tadpole / natural_spawn blastula | 6.69x10^-13^ | 2.22x10^-16^ | Inf | NT_Prick_UV neurula / NT_UV neurula | 1.88E-10 | 2.22x10^-16^ | Inf |
| NT_UV tadpole / NT_Prick blastula | 1.94x10^-10^ | 2.22x10^-16^ | Inf | NT_Prick_UV neurula / natural_spawn tadpole | 1.99E-14 | 2.22x10^-16^ | Inf |
| NT_UV tadpole / NT_Prick_UV blastula | 1.13x10^-09^ | 2.22x10^-16^ | Inf | NT_Prick_UV neurula / NT_Prick tadpole | 0.04118 | 2.22x10^-16^ | Inf |
| NT_UV tadpole / NT_UV blastula | 1.39x10^-09^ | 2.22x10^-16^ | Inf | NT_Prick_UV neurula / NT_Prick_UV tadpole | 0.4407 | 2.22x10^-16^ | Inf |
| NT_UV tadpole / natural_spawn gastrula | 6.69x10^-13^ | 2.22x10^-16^ | Inf | NT_Prick_UV neurula / NT_UV tadpole | 0.01562 | 2.22x10^-16^ | Inf |
| NT_UV tadpole / NT_Prick gastrula | 161.1 | 2.22x10^-16^ | Inf | NT_UV neurula / natural_spawn tadpole | 1.055x^-04^* | 1.08x^-05^ | 0.001033 |
| NT_UV tadpole / NT_Prick_UV gastrula | 3.37x10^-09^ | 2.22x10^-16^ | Inf | NT_UV neurula / NT_Prick tadpole | 2.19E+08 | 2.22x10^-16^ | Inf |
| NT_UV tadpole / NT_UV gastrula | 2.47x10^-09^ | 2.22x10^-16^ | Inf | NT_UV neurula / NT_Prick_UV tadpole | 2.34E+09 | 2.22x10^-16^ | Inf |
| NT_UV tadpole / natural_spawn neurula | 6.69x10^-13^ | 2.22x10^-16^ | Inf | NT_UV neurula / NT_UV tadpole | 82900000 | 2.22x10^-16^ | Inf |
| NT_UV tadpole / NT_Prick neurula | 13.83 | 2.22x10^-16^ | Inf | natural_spawn tadpole / NT_Prick tadpole | 2.07x10^12^ | 2.22x10^-16^ | Inf |
| NT_UV tadpole / NT_Prick_UV neurula | 64.02 | 2.22x10^-16^ | Inf | natural_spawn tadpole / NT_Prick_UV tadpole | 2.22x10^13^ | 2.22x10^-16^ | Inf |
| NT_UV tadpole / NT_UV neurula | 1.21x10^-08^ | 2.22x10^-16^ | Inf | natural_spawn tadpole / NT_UV tadpole | 7.86x10^11^ | 2.22x10^-16^ | Inf |
| NT_UV tadpole / natural_spawn tadpole | 1.27x10^-12^ | 2.22x10^-16^ | Inf | NT_Prick tadpole / NT_Prick_UV tadpole | 10.7 | 2.22x10^-16^ | Inf |
| NT_UV tadpole / NT_Prick tadpole | 2.636 | 2.22x10^-16^ | Inf | NT_Prick tadpole / NT_UV tadpole | 0.3793 | 2.22x10^-16^ | Inf |
| NT_UV tadpole / NT_Prick_UV tadpole | 28.21 | 2.22x10^-16^ | Inf | NT_Prick_UV tadpole / NT_UV tadpole | 0.03545 | 2.22x10^-16^ | Inf |

^1^Lower confidence limit, ^2^Upper confidence limit, * Significant pairwise comparisons, ^3^Inf=infinity
